# Spatial motifs reveal patterns in cellular architecture of complex tissues

**DOI:** 10.1101/2024.04.08.588586

**Authors:** Zainalabedin Samadi, Amjad Askary

**Author notes:** Contributing authors.

## Abstract

Spatial organization of cells is crucial to both proper physiological function of tissues and pathological conditions like cancer. Recent advances in spatial transcriptomics have enabled joint profiling of gene expression and spatial context of the cells. The outcome is an information rich map of the tissue where individual cells, or small regions, can be labeled based on their gene expression state. While spatial transcriptomics excels in its capacity to profile numerous genes within the same sample, most existing methods for analysis of spatial data only examine distribution of one or two labels at a time. These approaches overlook the potential for identifying higher-order associations between cell types – associations that can play a pivotal role in understanding development and function of complex tissues. In this context, we introduce a novel method for detecting motifs in spatial neighborhood graphs. Each motif represents a spatial arrangement of cell types that occurs in the tissue more frequently than expected by chance. To identify spatial motifs, we developed an algorithm for uniform sampling of paths from neighborhood graphs and combined it with a motif finding algorithm on graphs inspired by previous methods for finding motifs in DNA sequences. Using synthetic data with known ground truth, we show that our method can identify spatial motifs with high accuracy and sensitivity. Applied to spatial maps of mouse retinal bipolar cells and hypothalamic preoptic region, our method reveals previously unrecognized patterns in cell type arrangements. In some cases, cells within these spatial patterns differ in their gene expression from other cells of the same type, providing insights into the functional significance of the spatial motifs. These results suggest that our method can illuminate the substantial complexity of neural tissues, provide novel insight even in well studied models, and generate experimentally testable hypotheses.

## 1 Introduction

A central theme in biology is the idea that function is shaped by structure. Biological tissues, for example, often comprise stereotyped organizations of specific cell types that together enable proper function of the tissue. Formation of these structures during development is orchestrated by intrinsic gene regulatory networks and extrinsic cell-cell interactions. Therefore, analysis of cellular architecture of tissues can provide insight into both developmental processes that generate them and mechanisms that govern their function. Recent technological developments in spatial transcriptomics have enabled researchers to map the position of cell types in complex tissues, such as retina and brain [1, 2]. Developing methods to analyze and interpret the resulting cellular maps is an active area of research [3]. Reported developments can roughly be classified into two groups: methods that examine spatial distribution of one cell type label relative to itself and those that analyze the relationship between pairs of labels. Ripley’s spatial statistics [4] and spatial autocorrelation [5] are examples of the former. They determine whether cells with a given label are clustered, dispersed, or randomly distributed in space. The latter category includes methods that quantify association between label pairs based on proximity or predict activity of signaling pathways based on local expression of their ligands and receptors [3].

The important advantage of spatial transcriptomics lies in its multiplexity. Traditional RNA in situ hybridization and immunostaining methods could only map the expression of a few genes in one sample. In contrast, new spatial techniques can profile the expression of many genes, or even the entire transcriptome [6]. In this way, the position of many cell types can be mapped in the same sample. However, this feature is not directly utilized in virtually any of the existing analysis methods, as they focus on distribution of one or two variables at a time. Higher order associations between genes or cell type distributions are therefore not explored. Complex tissues, like those of the central nervous system, have highly organized structures made of many functionally distinct cell types. Therefore, quantitative analysis of spatial patterns that involve more than two cell types may be essential for understanding their development and function.

Spatial omics data are often modeled as neighborhood graphs. Graph theory offers a variety of methods for characterizing the topological and geometric organization of complex networks. Properties like degree distribution [7], diameter and average distance [8], clustering coefficient [9], and centrality measures [10] are commonly used to reveal global features of networks. However, these measures alone do not provide sufficient insight into the relationship between a network’s structure and its function. To better understand this relationship, it is necessary to study local structures of the network. Network motif analysis provides a framework for this, through identifying overrepresented subnetworks that may play a role in the network’s function [11]. This approach has been remarkably successful in identifying building blocks of biological networks, such as regulatory feedback and feed-forward loops. However, in neighborhood graphs, unlike genetic interaction networks or neural networks, the edges do not encode a functional relationship. The edge structure of a neighborhood graph depends to a large extent on the choice of the method for building the graph from node coordinates. In contrast, the node labels, which represent essential cell type identity information in spatial data, have been of less concern in previous work on network motifs. Therefore, existing methods for network motif analysis that focus on patterns of interactions between the nodes, rather than the identity of the nodes themselves, are not suitable for analysis of spatial data.

Here, we propose a strategy for identifying “Spatial Motifs” by uniform sampling of neighborhood graphs. Our approach extends the concept of motifs to spatial maps of cell types and identifies statistically overrepresented spatial arrangements in complex tissues. To achieve this goal, we developed an algorithm for enumeration and uniform sampling of paths in neighborhood graphs. Each path consists of a sequence of nodes, labeled by the cell types they represent. Paths along which the physical distance monotonically increases capture arrangements of cell types in the sample. We then adapted the STREME algorithm, originally developed to discover motifs in nucleic acid sequences, to identify overrepresented sequences of cell types in the sampled paths (Fig. 1). Our new algorithm, called Spatial MOtif REcognition (SMORE), takes into account that its input samples are derived from graphs, not one-dimensional DNA sequences. We tested sensitivity, specificity, and accuracy of SMORE by recovering motifs that were embedded at specified frequencies within synthetic graphs. We then analyzed published datasets of retinal bipolar cells and mouse hypothalamic preoptic region and identified spatial motifs that include a variety of neuronal and non-neuronal cell types. Our results also revealed that gene expression of cells in some spatial arrangements can differ significantly from other cells of the same type. Together, this work presents a novel and broadly applicable approach for identifying patterns in spatial data that go beyond pairwise associations.

**Figure 1:**
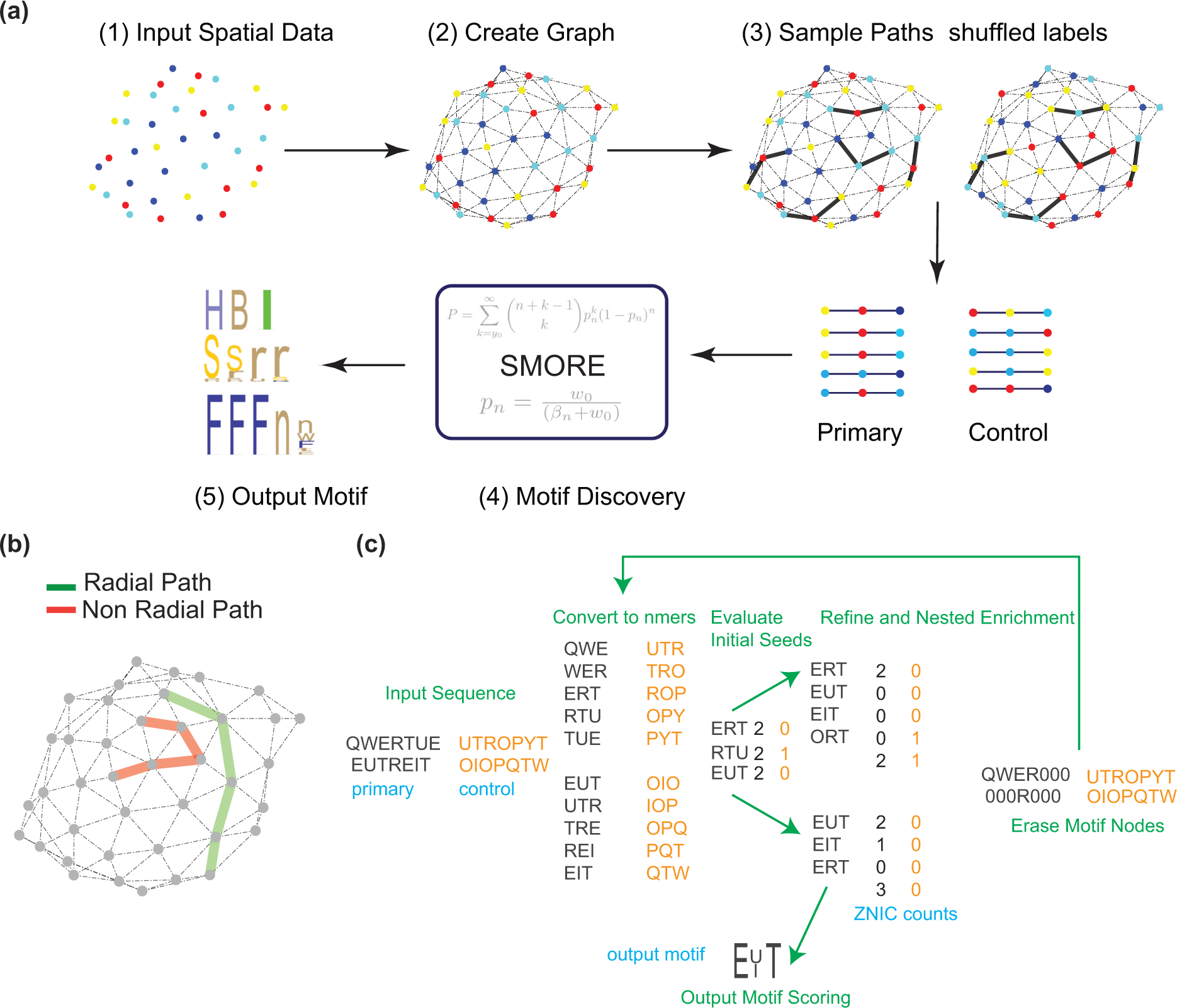
Overview of the spatial motif discovery algorithm. **(a)** schematic of the procedure for finding motifs on a set of spatially distributed nodes which are labeled by a set of colors or alphabets. (1) set of 2D distributed nodes (2) Delaunay triangularization is used in this case to generate the graph (3) URPEN is used to sample a set of 5 length-3 radial paths. Control data is created by shuffling the observed graph node cell types. (4) SMORE is used to extract most significant spatial motifs (5) PWM logos can be used to represent output motifs of different lengths **(b)** an example of a radial and a non-radial path. Radial path sampling ensures nodes that are farther from each other in the sample are also farther from each other in physical space. **(c)** A simplified example for the SMORE pipeline. Input samples are converted to nmers of the length of the motif and pvalues of the seeds evaluated based on negative binomial test with Bernoulli probability computed based on total number of distinct seeds in primary and control graph. nEval = 3 most significant seeds are input to refine and enrichment where ZNIC pvalues are used to refine and enrich candidate seeds. Hold-out scoring is performed to compute the pvalue for the output motif. Nodes of the seeds involved in the motif are erased from the graph and the process is repeated again. More detailed description of each block is provided in the methods section.

### 2 Design of the spatial motif discovery algorithm

Our algorithm for finding spatial motifs in neighborhood graphs consists of two main components; first a method to uniformly sample paths from the graph, and second, a procedure to find motifs in the obtained samples. Each path provides a sequence of cell type labels that occur near each other in space. Generating sequence samples from the neighborhood graphs reduces identification of spatial patterns into finding overrepresented sequences of labels. Despite some important differences, this task is similar to identifying motifs in nucleic acid or protein sequences. Therefore, we generalized existing methods of motif discovery in genomic sequences for our application on graphs.

The sampling algorithm takes a graph *G* and returns an unbiased sample of all paths inside the graph. In a graph, a path is a walk that does not intersect itself. Our selected paths are also constrained to be “radial”. Radial condition in a spatially-embedded network is defined as the requirement that physical distance along a path monotonically increases along the sequence of edges in the path. Radial condition ensures that the sequence of labels in a sampled path corresponds to a spatial arrangement of cell types in space, so that labels that are farther from each other in the path are also farther from each other in physical distance (Fig. 1b). Therefore, the radial requirement simplifies interpretation of the output motifs.

After sampling, the motif discovery algorithm identifies sequences of a given length that are statistically overrepresented in an iterative process. In each step, a significant recurring pattern of cell types is identified and is subsequently refined by considering sequences similar to the initial pattern or seed. The algorithms for path sampling and motif discovery are described below. For more details, refer to the extended methods section.

### 2.1 Sampling the graph to generate paths

The first step in our approach involves uniform sampling of neighborhood graphs. We have developed an algorithm for Uniform Random Path Enumeration (URPEN) based on the Rand-ESU algorithm [12]. The Rand-ESU method involves enumerating all potential subgraphs within a given graph, incorporating a probability element to uniformly sample a subset of these subgraphs. In our modification, we have adapted this method to exclusively sample paths as opposed to subgraphs. Paths differ from subgraphs in that they cannot intersect themselves, and each node, excluding the initial and terminal nodes, is only linked to its preceding and succeeding nodes in the path sequence. This contrasts with subgraphs where nodes can be connected to an arbitrary number of neighboring nodes. This distinction is crucial when selecting the next neighbor to expand the growing sample.

#### 2.1.1 Enumerating All length-k Paths in a Graph

The Path Enumeration algorithm (PEN), (Algorithm 1), enumerates all paths of length *k* within a graph. The algorithm begins with a vertex *v* from the input graph and adds only those vertices to the set that are neighboring the newly added vertex *w* but are not already in *V_path_*. To prevent enumeration of both the path and its reverse, the index of the last vertex in the enumerated path must be greater than that of *v*, though this requirement is not necessary for directed graphs.

#### 2.1.2 Uniform Path Sampling

Similar to ESU-tree, the PEN algorithm’s structure can be visualized as a tree structure. The tree structure for an example graph is demonstrated in (Fig. 2a). This tree has 18 leaves which correspond to the 18 size-3 paths of the graph. We can use this tree to sample paths uniformly without bias. The PEN algorithm systematically traverses its associated PEN-tree. In situations where a full traversal is impractical, we can perform a partial exploration of the PEN-tree such that each leaf is reached with equal probability. To achieve this, a probability is introduced for each depth 1 *≤ d ≤ k* in the path (or each depth in the PEN-tree), and the subsequent node rooted at a node at depth d is traversed with probability *p_d_*. This is implemented by calling the ExtendPath function at lines 3, and E6 of the PEN algorithm (Algorithm 1) with probability *p_d_*. This new algorithm is called Uniform Random Path Enumeration, URPEN. It can be observed that URPEN visits each path with equal probability of *p* = ^IT^*^k^ p_d_*. The method is tested in section 3.1 on a random graph to validate its accuracy numerically.

**Figure 2:**
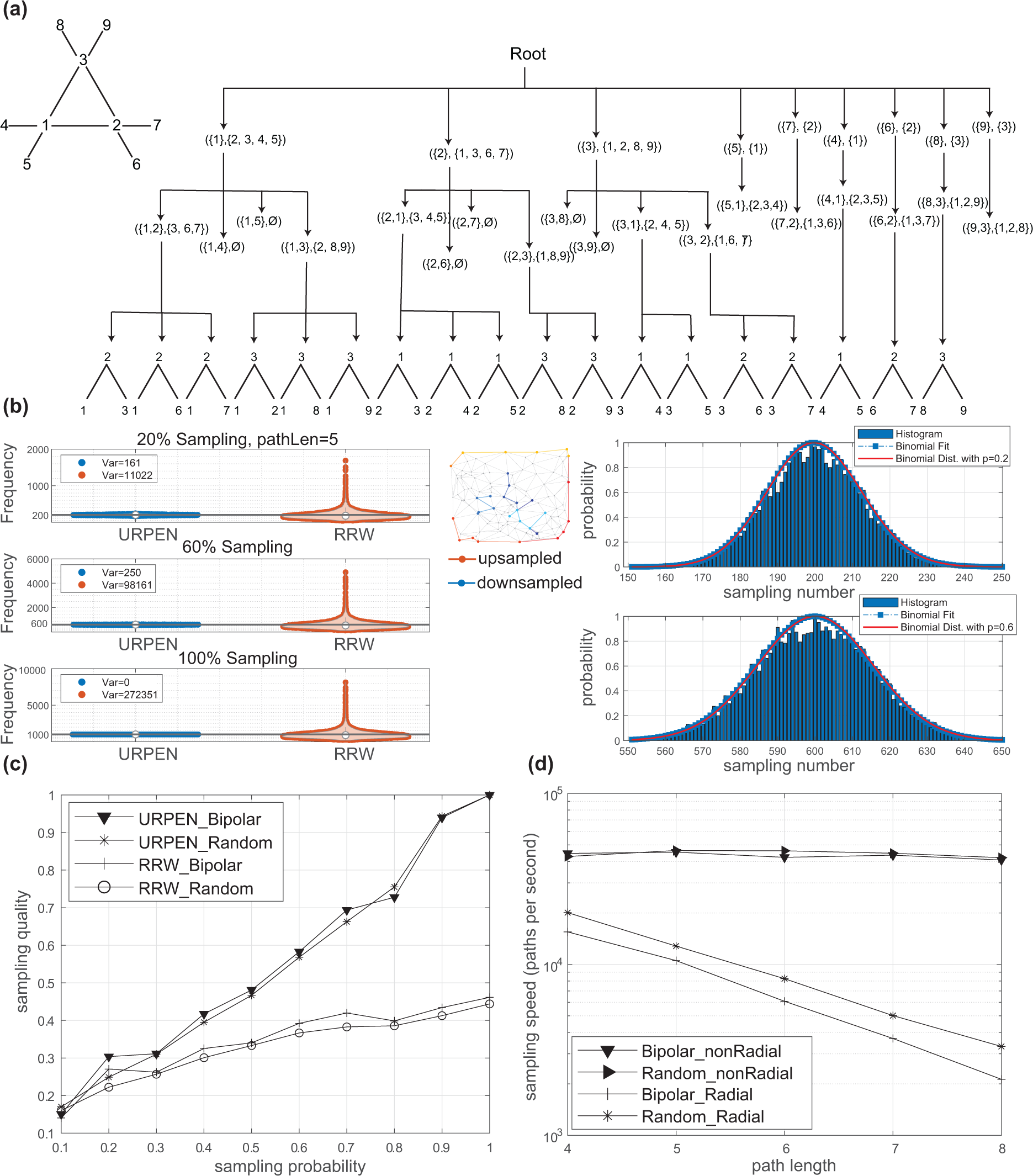
Unbiased sampling of paths from neighborhood graphs. **(a)** An example of the PEN sampling tree. **(b)** URPEN returns each path at a frequency corresponding with the sampling level, whereas Radial Random Walk (RRW) results in biased sampling of the graph. Examples of RRW upsampled and downsampled paths are shown in the top right corner of the 20% sampling panel. The sampling probability in URPEN is set to p = (1, 1, . . ., 1, 0.2) for 20% sampling, and p = (1, 1, . . ., 1, 0.6) for 60% sampling. This test was performed 1000 times. **(c)** Sampling quality with URPEN and RRW on the bipolar cell type graph and a random graph with 12000 nodes. **(d)** sampling speed for non-radial and radial sampling on the bipolar and random networks.

### 2.2 Spatial MOtif REcognition (SMORE)

Applied to a neighborhood graph, URPEN returns sequences of cell types that are observed near each other. Similar to DNA sequences, we can search through these cell type sequences for motifs using unsupervised learning techniques. The SMORE method is developed to detect motifs within the sampled sequences. Our approach uses as its basis the recently developed STREME method [13] which authors have demonstrated to be more accurate, sensitive, and comprehensive than several widely used motif discovery algorithms. STREME is developed to find motifs within sequence-like samples, SMORE on the other hand finds motifs in a network-based dataset. The algorithm follows a series of steps to accomplish its objectives.

1. Construct the graph from spatial data: To construct a graph from the spatial coordinates of cell types, Delaunay triangularization is employed, forming a graph with nodes as cells, labeled with their respective cell types. Given that each dataset may comprise multiple tissue sections or animal IDs, separate graphs are generated for distinct sections and IDs. For generating control data, each section or animal ID is shuffled independently. Additionally, besides Delaunay triangularization, the method offers options to construct the graph using arbitrary *K* nearest neighbors or epsilon graph approaches.
2. Sampling the graph and generating control data: SMORE uses URPEN for uniform sampling of the input graph. Control samples are generated using one of two methods: “shuffle” or “kernel”. The “shuffle” method produces control data by shuffling node cell type labels within samples (e.g., tissue sections and animal IDs). On the other hand, the “kernel” method establishes a kernel around each cell and swaps the cell’s label with that of a cell within its kernel (Fig. S1). In the experiments detailed in this paper, kernels with a radius of *K* neighbors are used, where *K* is specified for each experiment. *K* = 1 means only first neighbors within the graph are considered for shuffling. The degree of randomness in the control data is controlled by adjusting the number of neighbors considered for shuffling. There are certain scenarios where specific cell types associated motifs are readily apparent. In such cases, these cells can be fixed between primary and control data, meaning that their cell labels are not shuffled. We generate nTrain control data and seed numbers for control data are the total number of that specific seed within these nTrain samples. NScore independent control data is generated for output motif scoring, with the same settings as the training data.
3. Convert to n-mers and count seeds: Input samples to the algorithm can be of an arbitrary length. These samples are converted to W-mers, where W is the desired motif length. These W-mers are input to the count seed modules, where SMORE assumes that motifs don’t have any node in common (Zero Node in Common model; ZNIC). ZNIC counts of the unique seeds are computed, and along with their respective nodes are passed to the next module of evaluating initial seeds.
4. Initial Seed Evaluation: The significance of each initial seed is obtained by the negative binomial test. The justification for this specific test is argued in the extended methods section. The first nEval seeds with this criterion are passed to the next stage of refinement and enrichment. The default value for nEval is 25.
5. Refinement and Nested Seed Enrichment: The refinement and seed enrichment both use the same process of enrichment, except that refinement is only one iteration, and seed enrichment is nEnrich iterations, with the default value of nEnrich = 20. Enrichment groups similar path samples together and compares ZNIC counts of the grouped sequences with the control data. nEval motifs from the initial evaluation step are first enriched for one iteration and top nRef (nRef=4 as a default) motifs are further refined in seed enrichment block for nEnrich iterations or until pvalue doesn’t improve. During each iteration, the Position Weight Matrix (PWM) is calculated from the sequences participating in the motif. Likelihood ratio scores are then computed for all samples, using this PWM matrix, and the samples are arranged in descending order based on their PWM scores. In the event of an equal PWM score, the samples are further sorted based on their pvalues obtained in the initial evaluation block. Subsequently, ZNIC counts for the ordered samples are determined, and the PWM score threshold that minimizes the pvalue is identified. This process is iterated if the pvalue obtained is more significant than the previous iteration.
6. Motif scoring: In most cases, tissue samples are different from each other and it is not optimal to take sections of the samples as hold-out for scoring. In our experiment, the same samples are used for finding motifs and scoring the output motif, with the scoring part iterated over nScore times with different shuffled networks to avoid false positives. Implementation results on synthetic data with nScore = 50 shows that the false positive rate is negligible. Further, tests on real data with randomly shuffled cell type labels did not result in any significant output motif. The ZNIC counts for the seeds involved in the output motif are computed in the primary and nScore randomly generated control data and the 95 percentile least significant pvalue is considered as the output pvalue of the scoring module.
7. Motif node erasing: The respective nodes for the seeds involved in the motif are erased (i.e., their cell type is set to 0) from the graph in primary and control data along with their reverses. The previous steps are then repeated until output motifs are not significant anymore, or a specified number of output motifs have been discovered.

## 3 Results

### 3.1 URPEN enables efficient and unbiased sampling of neighborhood graphs

The sequence of cell types along a path in the neighborhood graph captures their local spatial arrangement. A collection of these sequences can be used for identifying overrepresented patterns in the graph, only if it represents an unbiased sample of all possible paths. Further, the sampling algorithm should be able to handle the large number of cells in typical spatial transcriptomics datasets.

To confirm that URPEN sampling is unbiased, we generated a graph by applying Delaunay triangularization on a spatially uniform distribution of 120 two-dimensional Cartesian points. We then used URPEN to sample radial paths of length 5 from this random graph at three sampling levels; 20%, 60%, and 100%. If sampling is unbiased, we expect each path to appear in the sample with a probability equal to the sampling level. We repeated these tests 1000 times. So, for the case of 20% sampling, we expect each path to appear on average 200 times in the output results. This number is 600 for 60% sampling and 1000 for complete 100% sampling. As a comparison, we also sampled the same graph with the commonly used random walk method. Random walk starts with a randomly chosen node and subsequent nodes are selected from the neighboring nodes with equal probability, until a path of the desired length is obtained. While URPEN returned paths at the expected frequency, random walk sampling showed significant bias for certain paths (Fig. 2b). The distribution of the URPEN counts is also consistent with a set of identical independent binomial distributions with *p* = *p_d_*, confirming that paths were sampled with equal probability (Fig. 2b). Similar results were obtained for other path lengths and for paths not constrained by the radial condition (Fig. S2). The sampling bias of random walk can be mitigated by using unbiased estimators [14]. However, that incurs complexity and leads to a penalty in speed performance [12].

We also evaluated the sampling quality of URPEN compared to random walk (Fig. 2c). Sampling quality is defined as the percentage of path types for which the number of extracted paths has at most 20 percent error relative to the exact counts, similar to the measure used previously [12]. Path samples with at least 5 counts were considered for quality evaluation. Sampling quality was computed for two cases: A random graph with 12000 nodes and 35823 edges and a graph based on spatial distribution of bipolar cells in a section of mouse retina [15]. Sampling quality for the URPEN method increases with increasing sampling probability, reaching one at complete sampling. In contrast, sampling quality for the random walk peaks at around 0.5.

We then assessed how sampling speed scales with the path length (Fig. 2d). The two graphs of Fig. 2c was sampled by URPEN at 10% level. The speed appeared to be independent of the path length for non-radial paths. However, when the radial constraint was applied, the speed decreased with path length, because the proportion of paths that meet the radial condition decreases with increasing path length. Consistent with this explanation, the speed of sampling radial paths is not independent of graph architecture, as it appears to be the case for the non-radial paths. Together, our results demonstrate that URPEN avoids the shortcomings of random walk sampling and offers a robust method for uniform path sampling.

### 3.2 SMORE accurately identifies overrepresented cell type arrangements in synthetic graphs

Evaluating the performance of SMORE requires datasets with known ground truth. Since the presence and frequency of overrepresented spatial patterns in existing experimental data is unknown, we created synthetic data by embedding patterns within random graphs at known frequencies. Each graph has 12000 nodes, 35823 edges, and 12 cell types. The embedding percentage indicates the proportion of nodes used for pattern embedding. For example, 2% embedding indicates that 2% of the nodes (i.e., 240 nodes) were used for embedding patterns. In the case of a length 4 motif, this results in 60 sequence patterns. Other nodes in the graph were labeled randomly. Embedded patterns included one variable position, which was filled with one of two cell types with equal probability. Other positions in the pattern were assigned one predefined cell type. Although more complex patterns can also be considered, we chose our patterns so that they are sufficiently complex while still providing insight into the algorithm’s performance. The algorithm was run 100 times for each embedding frequency, generating 10 output motifs per run. The samples for length 4 motif included all possible radial paths inside the graph. For the length 5 motif, the graph was downsampled by URPEN and 65% of all radial paths were used.

The accuracy of a motif discovery algorithm describes its ability to recover accurate versions of overrepresented patterns [13]. We assessed the accuracy of SMORE by measuring the similarity of output motifs to embedded patterns (Fig. 3a). In each run, SMORE returns 10 motifs that are sorted based on their statistical significance. At embedding frequencies above 0.5%, the most significant output motif tends to be the one most highly correlated with the embedded pattern. At 1% and higher embedding frequency, the Pearson correlation coefficient between the extracted motifs and the embedded patterns was higher than 0.9 in all 100 tests. Based on these results, we expect SMORE to be able to accurately identify length 4 and 5 motifs even when they only occur at low frequencies in a sample.

**Figure 3:**
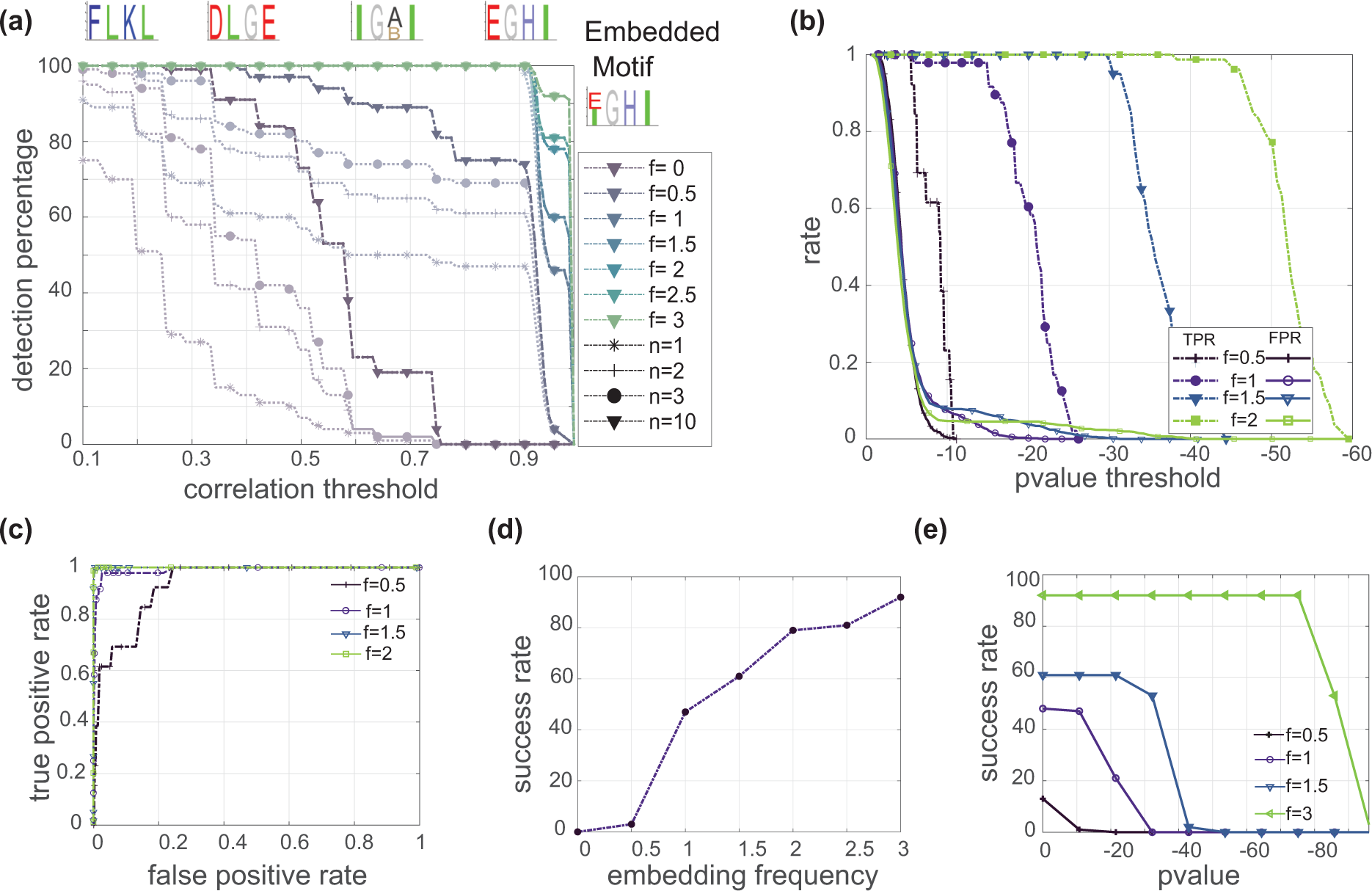
Evaluation of SMORE’s performance on synthetic data with known ground truth. **(a)** Accuracy of SMORE, defined as the best correlation coefficient between the embedded motif and extracted motifs for length 4 motifs. This figure represents the average results from 100 algorithm runs. The highlighted curves indicate the best correlation among the N = 10 output results. Faded curves demonstrate the results when considering only N = 1, 2, or 3 of the best output motifs, instead of all 10. Examples of motifs with different levels of correlation are shown on the top. **(b)** Specificity results for length 4 motifs, measured by false positive rate and true positive rates against log pvalue threshold and **(c)** against each other. **(d)** Sensitivity of SMORE measured by successful motif recovery rate against embedding frequency and **(e)** enrichment log pvalue. (See Fig. S3 for length 5 results)

To assess the specificity of SMORE, we examined the true positive rate (TPR) and false positive rate (FPR) against output pvalues (see Methods for details). Ideally, both TPR and FPR should be high at high pvalues and decrease as the pvalues decrease, with FPR decreasing at a faster rate to allow for correct motif identification. Accordingly, we observed that FPR curves reached zeros at around log pvalue of *−*10, while TPR performance gradually increases with the embedding frequency (Fig. 3b). The corresponding ROC curves (Fig. 3c) confirm close to perfect classification performance of SMORE at embedding frequencies higher than 1%.

To evaluate the sensitivity of SMORE, we defined success rate to be the proportion of output motifs that are statistically significant, at a given pvalue, and have a Pearson correlation coefficient of at least 0.95 with respect to the embedded pattern. With a log pvalue threshold of *−*10 (Fig. 3d), success rate for identifying length 5 motifs exceeds 90% at embedding frequencies above 1%. Success rates for length 4 motifs increase more gradually with embedding frequency, exceeding 90% only at 3% embedding. At each embedding frequency, success rate appeared to drop sharply beyond a specific pvalue threshold (Fig. 3e). As expected, this threshold decreases with increasing embedding frequency.

### 3.3 SMORE reveals spatial motifs in the distribution of mouse retinal bipolar cells

Retina contains more than a hundred neuronal cell types, organized into three layers of cell bodies. These cell types have different features and frequencies and are tiled across the retina in a stereotypical manner that supports the overall function of the tissue. Cell type diversity and individual variability make it difficult to identify recurring patterns in the cellular architecture of the retina. Spatial motif analysis can reveal higher order associations between retinal neurons and provide insight into their development and wiring.

We applied SMORE on a dataset of the mouse bipolar interneuron subtypes containing more than 30000 cells [16]. These subtypes were differentiated using co-detection of 16 gene markers by SABER-FISH, allowing the classification of all 15 bipolar subtypes. Bipolar interneurons bridge all visual circuits, establishing the link between sensory rod and cone photoreceptors and the output neurons. Bipolar cells also do not migrate from their birthplace, providing a spatial map between their final location and the location of their progenitors [17].

Rod Bipolar Cells (RBCs) constitute the majority of retinal bipolar cells in mice and their cell bodies are mainly organized together, further out in the Inner Nuclear Layer (INL) compared to Cone Bipolar Cells (CBCs) (Fig. 4a). SMORE evaluates the statistical significance of cell type arrangements in the experimental data against control data, which are generated by shuffling cell type labels. When shuffling is done for all cells across the whole graph, relatively obvious structures, like separation of RBCs, are identified as highly significant motifs (Fig. 4b, motif #2). To reveal other motifs involving RBCs, besides this trivial case, we can fix the position of RBCs in the control data (Fig. 4a). This would eliminate any motif whose significance stems from RBCs and elucidate the relationship between RBCs and other cell types (Fig. 4c, motifs #1 and 3).

**Figure 4:**
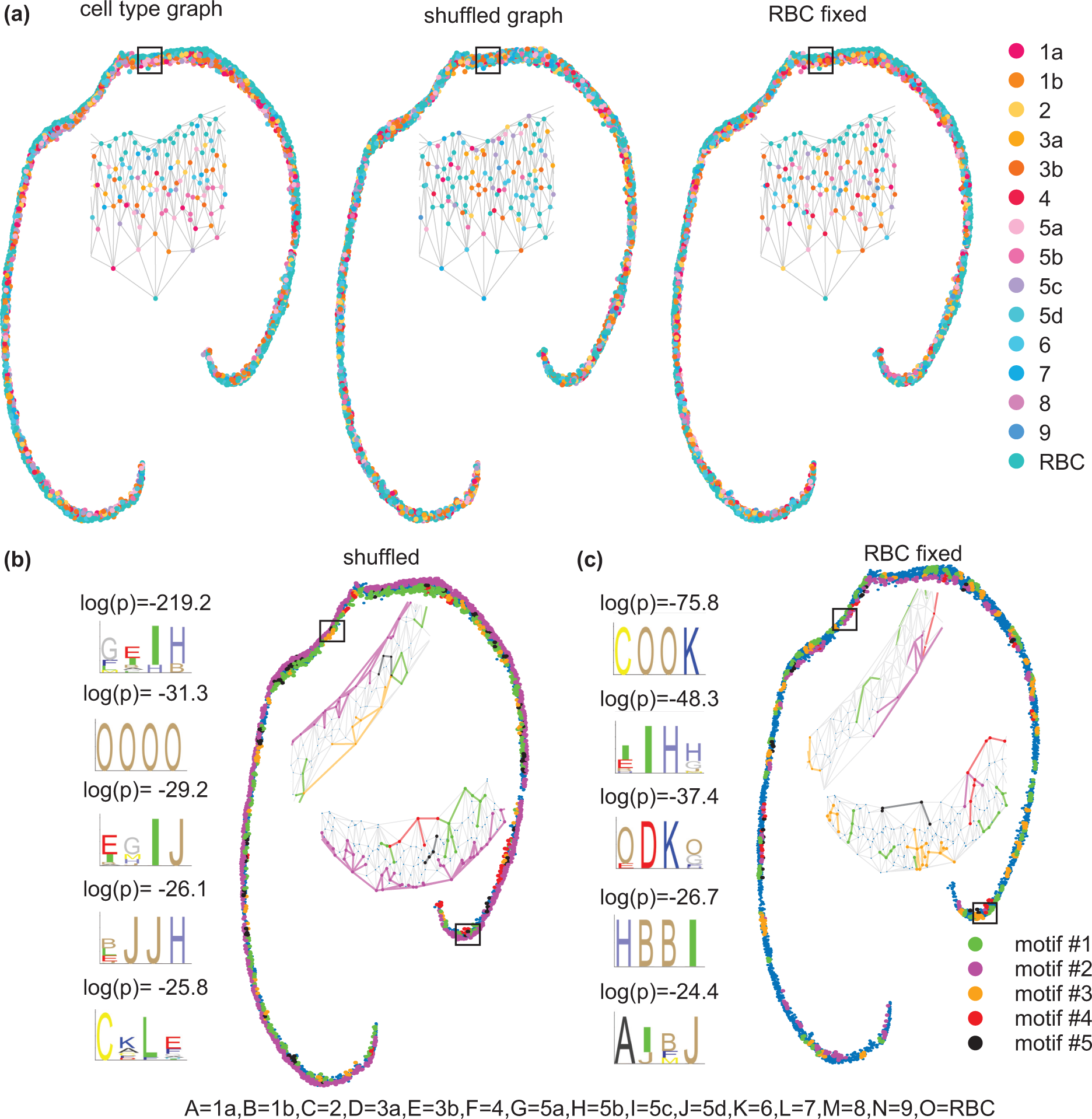
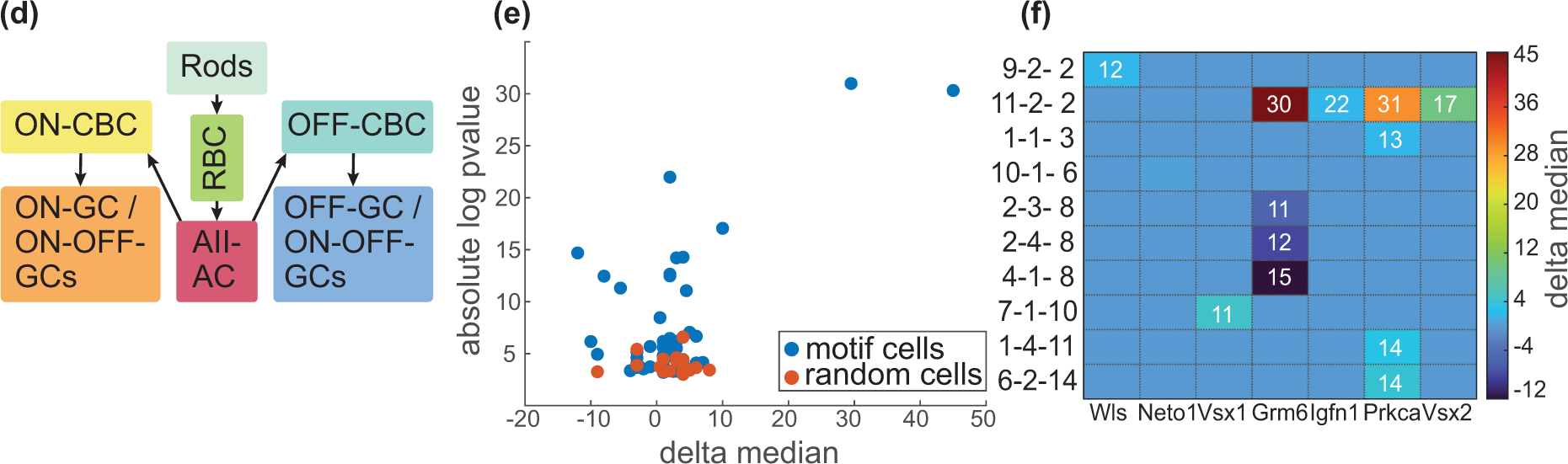
Spatial motif analysis of mouse retinal bipolar cells. **(a)** A retinal section with classified bipolar subtypes (left) and examples of control data generated from this section using different randomization methods, global shuffling (middle) and shuffling with fixed RBCs (right). **(b)** The first five output motifs, ordered from top to bottom, obtained using the global shuffling method of generating the control data along with their highlighted nodes on the tissue graph, and their respective log pvalues. **(c)** Same as **(b)**, with RBCs fixed. Annotations for the cell types involved in the motifs are listed at the bottom. **(d)** Schematic for primary pathway for rod-driven signals involving rods, rod bipolar cells, AII amacrine cells, OFF or ON (cone) bipolar cells, and OFF or ON ganglion cells. **(e)** Absolute log pvalue versus difference in gene expression medians (delta median) for cells in a spatial motif versus cells of the same type that are not in a motif arrangement. The results for a random selection of cells are shown in red. **(f)** Selected cases and genes with absolute log pvalue greater than 7. The heatmap is colored with delta median values. The motif cases, specified by “motif number-position-cell type”, are sorted by cell type on the vertical axis. Values in each cell show absolute log pvalue for comparison between cells within a given motif and overall cells of the same type.

Our analysis revealed several highly significant spatial motifs among retinal bipolar cells. These motifs can be investigated in the context of retina development, anatomy, and physiology. For example, when RBC positions are fixed, the most significant motif involves a type 2 OFF CBC followed by two RBCs and a type 6 ON CBC (“COOK” motif; Fig. 4c). Overrepresentation of this cellular arrangement can be understood in the context of a primary rod pathway that enables scotopic vision [18]. Within this pathway, the signal originating from a single rod cell is primarily directed to a select few AII amacrine cells through two RBCs [19]. The AII amacrine cells establish connections with nearly all bipolar cell types to gather scotopic signals originating from RBCs (denoted as O in our motif notation). These signals are then distributed to both ON and OFF CBCs through gap junctions and inhibitory synapses, respectively (Fig. 4d). However, the number of connections varies depending on the bipolar cell types [20]. Type 2 (C) cells account for 69% of the total number of OFF bipolar chemical synaptic contacts with AII amacrine cells, while type 6 (K) cells contribute 46% of the total area of ON bipolar gap junctions with AII amacrine cells [20]. Both type 2 and type 6 cells not only have the highest access to AII amacrine cell signals but also share these signals with other types of bipolar cells through interconnected gap junctions in the network. These findings support the central role of type 2 and type 6 cells in conveying the most sensitive scotopic signals to the postsynaptic ganglion cells. Further, AII amacrine cells are characterized by their narrow-field dendrites. Typically, a bipolar cell receives more inputs from AII amacrine cells that are in its close proximity [20]. This suggests that bipolar cell types involved in scotopic vision should be spatially close to each other. Given these considerations, we hypothesize that the “COOK” motif is associated with the primary rod pathway for scotopic vision in mice.

### 3.4 Cells within spatial motifs exhibit gene expression differences compared to other cells of the same type

Spatial context of the cells often influences their function. With recent advances in spatial gene expression profiling, there has been an increased interest in systematically characterizing spatial variability of gene expression [21–27]. Spatially variable genes may explain functional distinctions between cells in different regions or demarcate spatial domains [21–23, 26]. If spatial motifs represent functional units made of various cell types that together play a distinct role, we can expect that at least in some cases cells participating in these motifs exhibit distinct gene expression signatures.

We assessed differential gene expression between cells of each type that are involved in a spatial motif and the ones that are not. For each gene, the median expression value among the cells of a specific type that are involved in the motif was subtracted from the median of all the cells of the same type. The pvalues for the observed delta medians were obtained through theoretical computation by analyzing the distribution of delta median values for a random subset of cells (see Methods for details). We performed this analysis for motifs obtained by shuffling with fixed RBC cells for the 16 genes profiled in the retinal bipolar dataset [16]. Several cases of highly significant differential gene expression were observed in the motif cells (Fig. 4e). In contrast, control samples where a random subset of cells were selected, with the same size and type of the corresponding motif case, showed much higher pvalues. This observation is consistent with functional specialization of cells in spatial motifs.

The heatmap in Fig. 4e illustrates the absolute log pvalues across selected cases for 20 output motifs. Each motif can consist of multiple cell types in different positions. For example, the second motif in the fixed shuffling case of Fig. 4c comprises 10 cells, 4 cells in each of the positions 1 and 4, and one cell in positions 2 and 3. Here we consider each cell type in each position of each motif as a separate case. Genes with absolute log pvalues greater than 7 are highlighted in Fig. 4f.

Among genes that were significantly upregulated or downregulated in the motifs, Grm6 stands out because it shows the most extreme differential expression in both directions. Grm6 is upregulated in type 1b OFF bipolar cells (B) in OBBO motif, where O represents an RBC, (delta median = 45*, −* log(pvalue) = 30.31) and downregulated in type 5b ON bipolar cells (H) in “HBBI” motif (Delta median = *−*12*, −* log(pvalue) = 14.67). Grm6 encodes the metabotropic glutamate receptor 6 (mGluR6) which is localized to the dendritic tips of ON bipolar cells [28]. It plays a crucial role in triggering depolarization of ON bipolar cells in response to light-induced hyperpolarization of photoreceptors [29]. Mutations in Grm6 gene in humans lead to autosomal recessive congenital stationary night blindness (arCSNB) [30].

We observed that enrichment of Grm6 in type 1b cells in motif 11 (OBBO) can be explained by their position along the radial axis of the retina. Grm6 is expressed at higher levels in type 1b cells whose cell body is closer to the photoreceptor level (Fig. 5a). Since RBCs (O) are concentrated in this outer region, type 1b cells in “OBBO” motif also tend to be in radial positions where Grm6 expression is higher (Fig. 5a). In contrast, downregulation of Grm6 in type 5b cells (H) of motif 4 (“HBBI”) seems to be explained by their proximity to type 1b (B) cells rather than their radial position (Fig. 5b-d). Type 1b cells lack dendrites connecting them to photoreceptors [31, 32]. Therefore, the mechanism of their function is not well understood [31]. Our observation suggests that type 1b cells may influence signal processing in the retina by altering expression of key postsynaptic receptors, like Grm6, in nearby ON bipolar cells.

**Figure 5:**
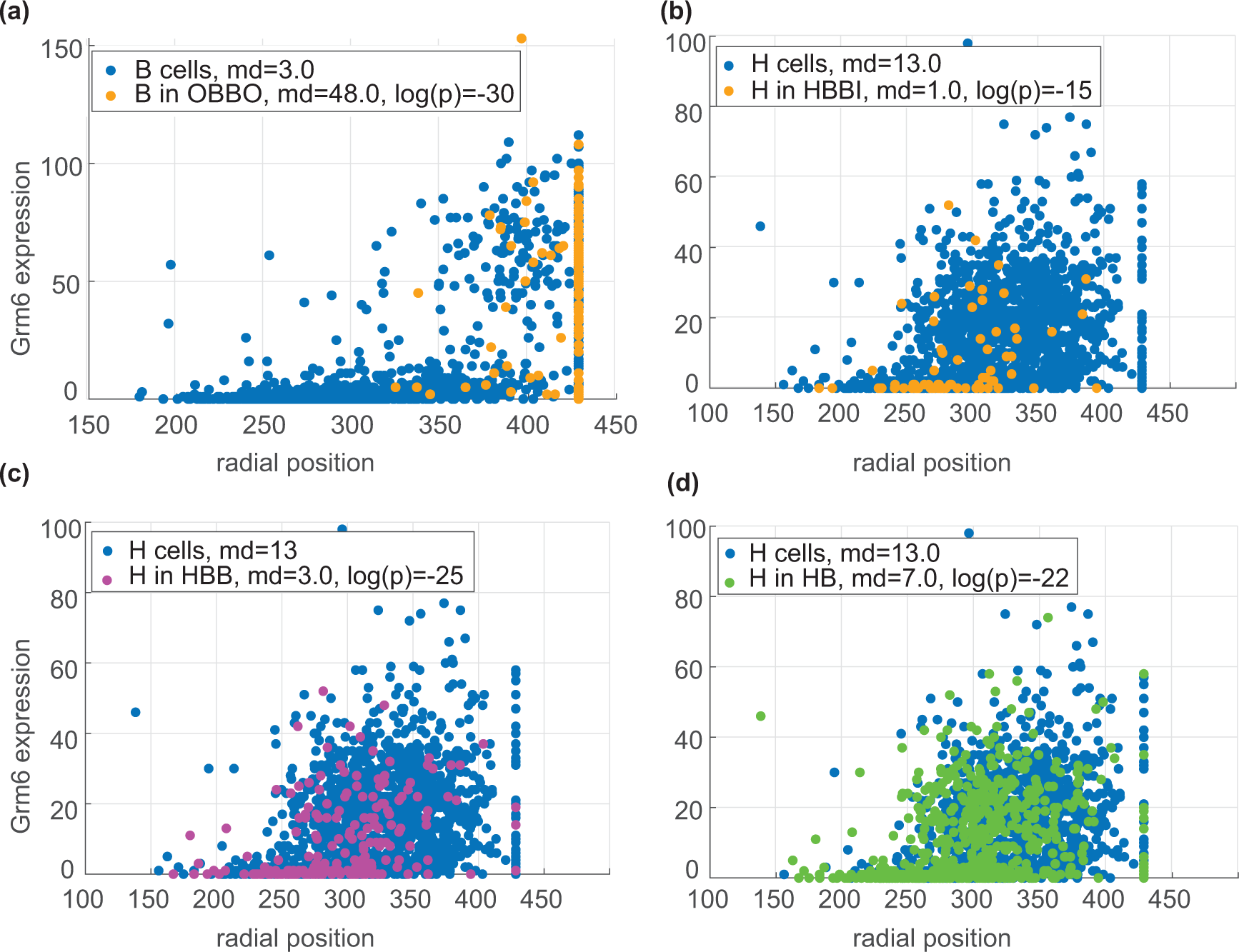
Grm6 shows motif specific expression patterns. Expression of Grm6 versus radial position of cell in the mouse retina for **(a)** type 1b and **(b-d)** 5b cells. Cells within specific spatial arrangements are highlighted in each panel. The radial position was computed by considering the outer extreme points as the maximum radius points and computing the radial position for other nodes relative to the nearest outer point.

### 3.5 SMORE reveals overrepresented cell type arrangements in the preoptic area of mouse hypothalamus

To explore the utility of SMORE at identifying non-trivial recurring patterns in a significantly more complex sample, we applied it to a spatial transcriptomics dataset of the mouse hypothalamic preoptic region [33]. This dataset profiles about 1 million cells and has identified about 70 neuronal populations with distinct signatures and spatial organizations. As we showed before, the approach used to generate control data affects the output motifs. This can be used to tune the algorithm to different anatomical features. Here in addition to fixing specific cell types, we introduce local kernel shuffling (Fig. 6a-c). Shuffling cell labels across the entire sample can result in the emergence of relatively straightforward structural motifs, such as regional boundaries. On the other hand, local shuffling maintains cell type frequencies within cellular neighborhoods. Therefore, if a sample is compartmentalized to regions with distinct cellular compositions, local shuffling is more likely to identify patterns that are overrepresented within each compartment. Control data generated by local shuffling maintains a higher degree of similarity with respect to the original experimental data. Therefore, the motifs obtained with global shuffling tend to have more significant pvalues compared to the output motifs of local shuffling. In both shuffling methods, if certain cell types form obvious structures (e.g., Fig. 6a Ependymal cells (blue)), their positions can be fixed in the control data.

**Figure 6:**
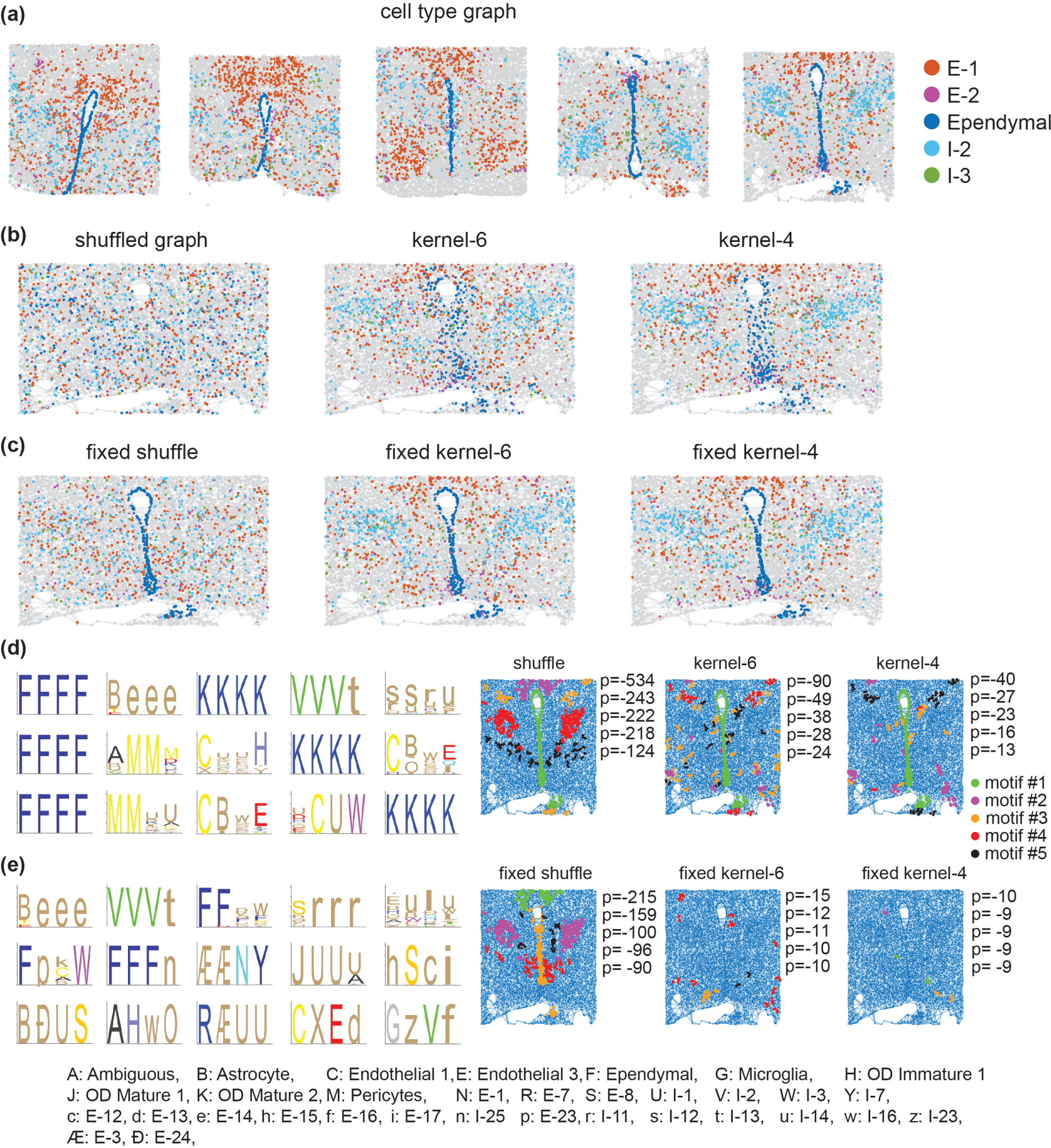
Spatial motif analysis of mouse hypothalamic preoptic region. **(a)** the graph of the preoptic region of the hypothalamus from five adult male mouse brains at Bregma −0.29. **(b)** Examples of control data generated from the 5th tissue using different randomization methods: global shuffling, kernel shuffling with 6 neighborhood depth, and kernel shuffling with 4 neighborhood depth. **(c)** control data obtained by applying the same randomization methods as (b) but with the positions of non-neuronal cell types fixed. **(d-e)** First five output motifs obtained using different methods of generating the control data along with their highlighted nodes on the tissue graph, and their respective log pvalues. Obtained motifs are represented by logos in the left panel, annotations for the cell types involved in the motifs are listed at the bottom. Motifs obtained using not fixed (d) and fixed non neuronal cell type (e) control data generation methods. Each motif is indicated by a different color in the highlighted tissue graph.

We applied SMORE to cellular maps of the hypothalamus preoptic region from five adult male mouse brains at Bregma*−*0.29 to identify length 4 spatial motifs. The sections used in our analysis comprise 35, 693 cells. We tried both global shuffling and local shuffling with kernel sizes of 4 and 6 (Fig. 6b). In another set of experiments, we also fixed the position of non-neuronal cell types between the primary and control data (Fig. 6c). When non-neuronal cell types were not fixed, the most significant motif consists of a group of four interconnected ependymal cells (Fig. 6d). This pattern is immediately visible in the graphs because ependymal cells form a layer that lines the ventricles. As expected, fixing the position of non-neuronal cells removes this motif as well as the motif made entirely of mature oligodendrocytes (Fig. 6e). Instead, other motifs emerge, some of which are combinations of ependymal and neuronal cells (e.g., motif 3 of fixed global shuffling).

The first five motifs generated through the global shuffling method correspond to discernible patterns within the fifth tissue section shown in Fig. 6a. In addition to the first motif of ependymal cells, motif 2 comprises astrocytes, E-9, and E-14 cell types, which are enriched in the PVA nuclei of the hypothalamus [33]. E-14 and E-9 cells also show similar gene expression patterns [33]. Motif 3 represents a pattern of mature oligodendrocytes known to be enriched in the anterior commissure and the fornix. Motif 4 is a pattern of I-2 and I-13 cells which are indicated to be enriched in BNST-p and StHy nuclei. I-2 and I-13 are both Aromatase-enriched clusters and express both androgen receptor (Ar) and estrogen receptor alpha (Esr1) [33]. Motif 5 is primarily composed of cell types I-11, I-12, and I-14, collectively enriched in the MPN and StHy nuclei.

Motifs obtained from kernel shuffling methods capture less obvious patterns. For instance, motif 5 in the case of “fixed kernel 6”, represented as hSci sequence which is equivalent to a radial path of 4 excitatory neuronal cell types, E-15, E-8, E-12, E-17, occurs 4 times in the primary data (2 occurrences in each of animal IDs, 10, and 11) and 3 times in the total of 50 generated control data. This motif consistently appears in kernel shuffled tests, with different ranks. Interestingly, there is a similar motif, BSic, with B representing astrocytes, that appears as motif 11 of the not fixed kernel 4 (Fig. S4a) experiment and appears at the opposite side of the brains of the same Animal IDs.

The functional explanation of identified spatial motifs is not the focus of this study. But it is reasonable to expect that such patterns hint to either functional relationships between the cell types involved or specific developmental programs that generate them. Therefore, they can help generate hypotheses for future studies. For instance, motif 5 in the non-fixed global shuffling experiment primarily consists of cell types I-11, I-12, and I-14. In the case of the fixed shuffle, motif 4 is predominantly composed of cell types E-8, E-15, I-34, and I-15 in the first position, while I-11 occupies the remaining positions. I-15, I-2, I-11, I-14, I-33, E-8, and E-15 cells display sexually dimorphic cFos enrichment in male mating [33]. Interestingly, a similar motif exists in female sections (Fig. S4b), where I-15 replaces I-11. The Esr1-enriched cluster I-15 exhibits significant enrichment in female animals and is preferentially activated in females, with lesser activation in males after mating [33].

We also performed gene expression analysis for the motifs obtained by global shuffling with fixed non-neuronal cells. Many cases of highly significant differential gene expression were observed in the motif cells (Fig. 7a). The heatmap in Fig. 7b illustrates the absolute log pvalues across all cases for 40 output motifs. The majority of statistically significant cases for genes imaged using combinatorial smFISH measurements are concentrated in the first 10 output motifs. In contrast, genes measured through sequential FISH rounds, typically genes with higher expression levels, exhibit a higher prevalence of significant cases in this analysis. This difference is probably related to the fact that gene expression values for sequential genes are generally under dispersed (Fig. S5), resulting in the upregulated or down regulated genes being more significant. Genes with absolute log pvalues greater than 20 are highlighted in Fig. 7c. In most cases, the significance of differential gene expression varies depending on the position in the motif. For example, Vgf is upregulated in motif 2, position 1, cell type 26 (i.e., 2-1-26 case). But its differential expression is not significant in position 3 of the same motif. This may indicate that within a spatial motif, cells of the same type in different positions can have differences in their gene expression and potentially distinct roles.

**Figure 7:**
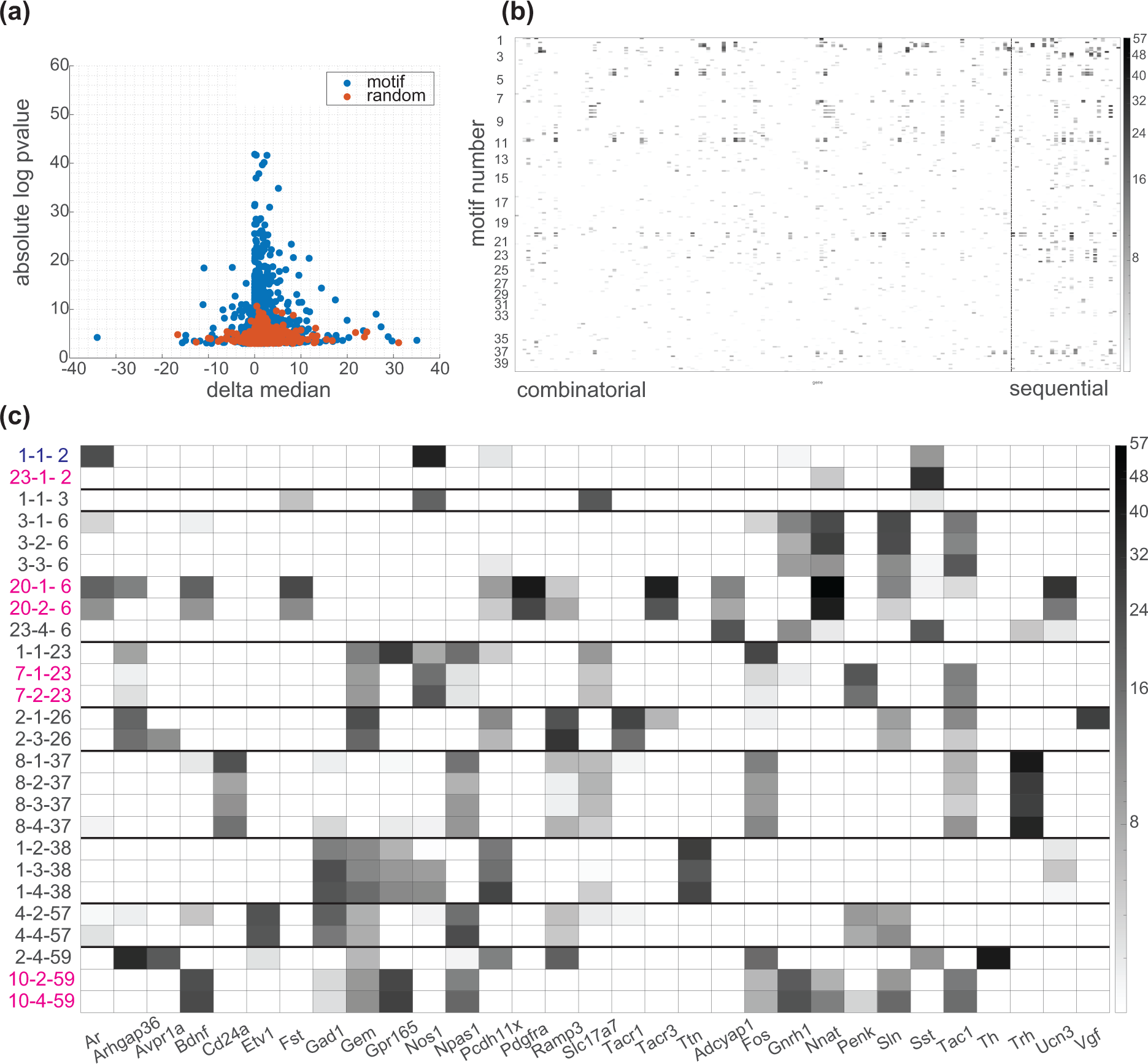
Motif specific gene expression analysis for hypothalamus motifs. **(a)** absolute log pvalue against difference in median gene expression in motif and non-motif cells. Blue markers show experimental results, red markers are the results for one random selection of cells. The expression values are not normalized. **(b)** heatmap of absolute log pvalues for genes (on the horizontal axis) and motif cases (on the vertical axis). Out of 155 genes, 135 were measured by combinatorial FISH and 20 were measured in sequential rounds of FISH. These two groups are separated with a dashed line. **(c)** selected cases and genes with at least one absolute log pvalue greater than 20. The motif cases are sorted by cell type. The gene names and their motif address (motif number-position-cell type) are included on the x- and y-axis, respectively.

### 3.6 Identifying spatial motifs in a 3D sample

So far spatial transcriptomics has mostly been applied to thin tissue sections or monolayer cultured cells. In these cases, obtained measurements are typically projected on a two-dimensional plane. However, recent advances have made it possible to map gene expression in thicker tissue slices, resulting in 3D datasets [34–37]. This is an important step in development of spatial methods because it enables profiling of cells in their native context, which in many tissues of interest is inherently three-dimensional. Since our method operates on a neighborhood graph, it should be able to identify spatial motifs in 3D datasets as well. To test this, we applied SMORE on the cellular map of a 200 µm thick slice of mouse anterior hypothalamus, encompassing over 78000 cells [38]. The cells in this dataset are classified into 21 excitatory neuronal clusters, 26 inhibitory neuronal clusters, and 7 non-neuronal cell subclasses based on the expression of 156 genes (Fig. 8a).

**Figure 8:**
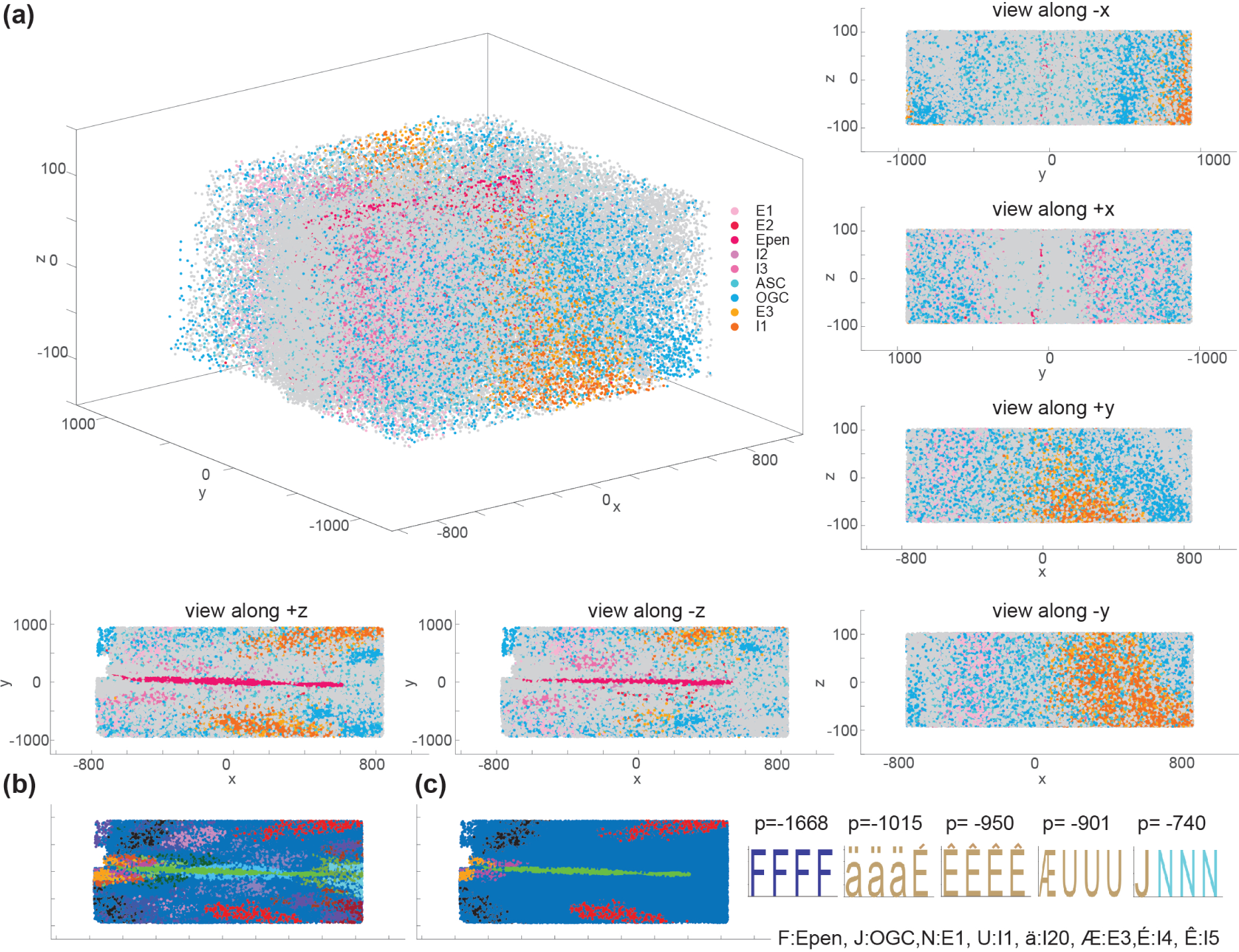
Spatial motif analysis of a 200 µm slice of mouse hypothalamus. **(a)** the graph of 3D MERFISH dataset with classified cell subtypes shown as dots colored by subtype. The perpendicular views from different angles are shown alongside the 3D view for the primary graph. Epen: Ependymal cells, ASC: astrocytes. OGC: Oligodendrocytes, Excitatory and inhibitory subtypes started with E and I letter, respectively. **(b)** the first 20 spatial motifs obtained using global shuffling to generate the control data. **(c)** the first 5 output motifs along with their highlighted nodes on the tissue graph, and their respective log pvalues. Annotations are the same as Fig. 6 for the shared cell types.

Fig. 8b illustrates the top 20 output motif nodes highlighted within the graph. Comparing this figure with the equivalent view from 8a (lower left) indicates the similarity of the highlighted motifs with the visual structure of tissue and agrees with our expectation that global shuffling is able to derive discernable patterns, along with other motifs that are less obvious. Fig. 8c highlights the first 5 motifs along with the logos representing the identified motifs. Overall, our results demonstrate that SMORE can be used for analysis of both 2D and 3D spatial data.

The cells within the 3D graph of tissue structure tend to have a greater number of neighbors on average compared to those in 2D datasets. For instance, while the average number of neighbors for the 2D mouse hypothalamus dataset in Fig. 6a is 5.9, it is 15.2 for the 3D hypothalamus dataset. Additionally, the 3D dataset comprises more than twice the number of cells in the 2D hypothalamus dataset (78, 229 compared to 35, 693). Consequently, the number of paths within the 3D dataset is substantially higher. Here we used URPEN to sample 10% of the radial paths in this 3D tissue graph. Despite this downsampling, the number of radial paths for the 3D dataset was 3, 676, 186 while it was 696, 141 for the specific tissue sections analyzed in Fig. 6, which were not downsampled. Due to the larger sample size, the log pvalues for the 3D dataset exhibit greater significance.

## 4 Discussion

Spatial transcriptomics is a rapidly growing field. We have seen significant innovation in the field over the past few years, resulting in a multitude of techniques and consistent improvements in their efficiency and scalability. Concurrently, application of spatial transcriptomics has expanded beyond specialized groups to the broader community of biomedical researchers. There are already several commercial platforms available to researchers, and it is expected that additional options will become available in the near future. Therefore, the need for innovative computational methods to extract biologically relevant information from this type of data is on the rise.

Methods of analysis of spatial transcriptomics data can be roughly divided into two groups. First, methods that analyze gene expression distribution in space. Combining spatial information with gene expression necessitates addressing spatial heterogeneity during preprocessing and clustering of gene expression [39–42]. Several methods focus on identifying regions of adjacent cells with similar gene expression [26, 42–47] or spatially variable genes with gradient, periodic, or spot-like patterns inside the tissue [21–27]. Spatially resolved gene expression can also be used to elucidate cell-cell interactions [48–50]. Interactions between cells are analyzed either by identifying interacting gene pairs [51–53] or identifying pairs of cells that express known interacting genes, such as ligand-receptor pairs [3, 42, 54]. Spatial domains associated with cell-cell communication can be also identified by clustering interacting cells downstream of communication inference [55]. Second group are methods that analyze spatial distribution of cell types. This includes methods for examining spatial distribution of each cell type by itself [4, 5] as well as pairwise enrichment analysis [56].

In this work, we introduce a method for identifying patterns of cell type arrangements with arbitrary length. There are two major contributions that make this task possible: a method for unbiased sampling of paths from a graph (URPEN) and a method for identifying motifs in such samples (SMORE). Our approach is general and can be applied to any system with sufficient complexity. This includes solid tumors and organoids, where greater heterogeneity increases the need for statistical analysis. It is also not limited to two-dimensional maps and can be readily adapted for three-dimensional data. In addition to the application presented here, individual components of our method can be independently employed in a range of other contexts that are modeled as a graph. For example, path sampling via random walk has been used in applications ranging from network representation learning [57] to estimation of similarity measures [58]. Unbiased path sampling using URPEN can offer benefits over random walk in such applications. Detailed evaluation of performance improvement with URPEN needs further investigation.

Spatial motifs can be explained in terms of their functional significance or developmental mechanisms that generate a specific cell type arrangement. Therefore, they can be used to generate hypotheses and further our understanding of tissue biology. We have provided a few examples in this study. This includes association of RBCs with type 2 and 6 cone bipolar cells that can be involved in the primary scotopic pathway and downregulation of Grm6 in type 5*b* ON bipolar cells near type 1b cells that offers a possible mechanism for the function of these atypical bipolar cells. Interpretation of each spatial motif at this point can only be done on a case-by-case basis, in the context of what is known about the cell types involved. This can begin with length 2 motifs, which are easier to interpret, and progress to higher lengths in a stepwise manner. For motifs that consist of more than one seed, each seed can also be investigated separately. It will be interesting to explore more systematic ways of utilizing spatial motifs for characterizing tissues. For example, the set of all spatial motifs in a sample might provide a quantitative representation of the tissue structure. These representations may prove valuable in classifying tissues with subtle differences in their cellular architecture, such as various cancer subtypes.

Our analysis here is constrained by certain technical limitations of the existing data. In datasets we analyzed, each cell is represented by a point in the tissue, which is typically the center of its nucleus. This ignores variation in size of the cells and their receptive fields. Position of the cell body may also not be a good indicator of neuronal connectivity. Further, our motifs are currently confined to short range local neighborhoods surrounding each cell. Expanding the method to incorporate longer range interactions could be an interesting next step, either by clustering identified motifs or by permitting gaps in the motif sequence. Gene expression analysis can be informative as shown here. However, gene panels in imaging based spatial transcriptomics datasets are often selective, focusing on known cell type markers. As more, scalable and multimodal spatial methods become available, SMORE has the potential to discover more intricate relationships between the spatial positioning of cells and their functional characteristics, including gene expression.

## Acknowledgements

We thank Swetha Ramesh, Roy Wollman, Eric Deeds, and Alexander Hoffmann for helpful discussion and feedback. This research was supported with funding from National Eye Institute of NIH under award number R00EY031782 and the UCLA Society of Hellman Fellows award.

## Ethics Declarations

### Ethics approval and consent to participate

Not applicable.

### Consent for publication

Not applicable.

### Availability of data and materials

The code for sampling and motif discovery algorithms is implemented in MATLAB and is publicly available under the MIT license at [59]. The dataset used for the mouse bipolar interneuron subtypes is publicly available at [60]. The dataset for mouse hypothalamic preoptic region is publicly available at [61], and 3D mouse anterior hypothalamus dataset is obtained from authors of [38] via personal communication.

## Supplementary Information

### Methods

In order to apply the SMORE method on the spatial structure of the cell types, spatial transcriptomics dataset is imported and the neighborhood graph based on cell positions is created. Control data is generated using one of two methods: global shuffling and kernel shuffling. As an example, Fig. S1 illustrates a kernel for the center node (highlighted in magenta). In this figure, *K* is set to 4 and 6, indicating that all nodes within *K* neighborhoods of the center node are part of the kernel.

nTrain instances of shuffled labels are generated. Labels for fixed nodes are not shuffled and labels for the other nodes are not shuffled with the fixed nodes. The graph is sampled with URPEN with the specified sampling frequency and the labels for the sampled paths are imported from either the original cell type labels or the set of nTrain shuffled labels created for the control data. After incorporating the reverse paths into the dataset, both primary and control data are fed into the SMORE method to identify motifs. The algorithms for path sampling and motif discovery are described in detail below.

### Uniform Path Sampling

The topological relations inside the spatial data can be represented as graphs. A graph is defined as an ordered pair *G* = (*V*; *E*) consisting of a nonempty set of vertices *V* and a set of edges *E* of two-element subsets of *V* . In the following, we’re dealing with undirected graphs, but the path sampling algorithm can equally be applied to directed graphs as well. Vertices in *V* are assumed to be uniquely indexed by the integers 1*, . . ., n*, where *n* = *|V |* is defined as the size of the graph. *v > u* is used to indicate that the index of a vertex *v* is larger than that of a vertex *u*. For a vertex *v* ∈ *V*_0_, where *V*_0_ is a subset of vertices, its forward neighborhood with respect to *V*_0_*, N_frw_*(*v, V*_0_), is defined as the set of all vertices from *V \ V*_0_ which are adjacent to *v*. The neighborhood of a vertex is simply its forward neighborhood with respect to the empty set, ∅.

The developed algorithm for finding motifs in graphs consists of two main components; first a method to uniformly sample the graph, and second, a procedure to find motifs inside obtained sequences. The sampling algorithm takes a graph *G* and all paths inside the graph are sampled uniformly. In a network, a path is a walk that does not intersect itself. Selected paths are also constrained to be radial. Radial condition in a spatially-embedded network is defined as the requirement that actual physical distance along a path monotonically increases along the sequence of edges in the path. The graph on spatial dataset is created based on the distance between nodes, therefore, the radial requirement enables us to better interpret output motifs in our experiment on the real dataset.

#### Algorithm 1

Path Enumeration (*G, k*)

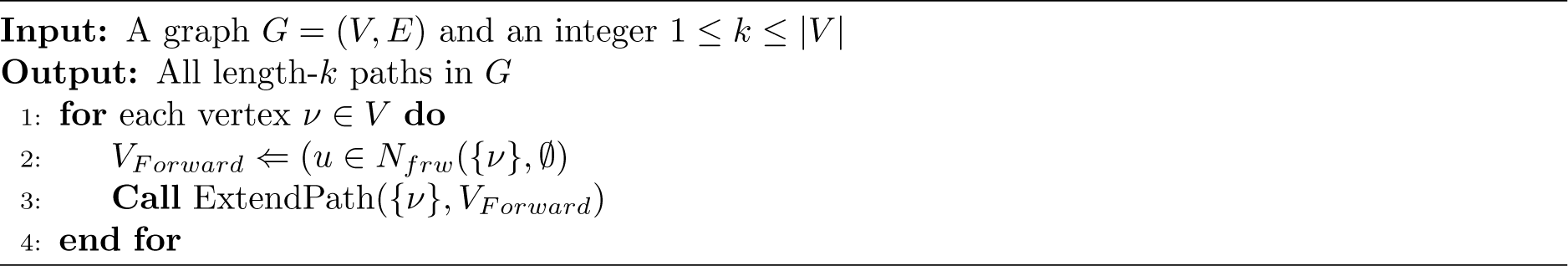

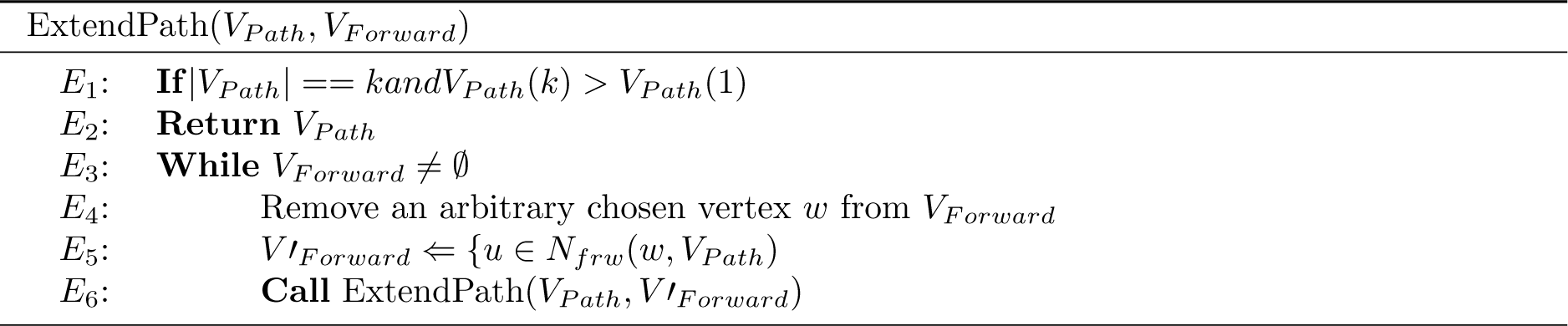

The Path Enumeration algorithm (PEN), (Algorithm 1), enumerates all paths of length *k* within the graph. The algorithm begins with a vertex *v* from the input graph and adds only those vertices to the set that are neighboring the newly added vertex *w* but are not already in *V_path_*. To prevent enumeration of both the path and its reverse, the index of the last vertex in the enumerated path must be greater than that of *v*, though this requirement is not necessary for directed graphs. The proof for the correctness of PEN is similar to that of the ESU algorithm [12].

We can modify the PEN algorithm to enumerate a subset of paths such that each path is reached with equal probability. This is implemented by calling the ExtendPath function at lines 3, and *E*_6_ of the PEN algorithm with probability *p_d_*. This new algorithm is called Uniform Random Path Enumeration, URPEN. The method is tested in section 3.2 on a random graph to validate its accuracy numerically.

### Compute Significance

The significance of each initial seed is obtained by the negative binomial test. The Poisson distribution arises naturally in the study of data taking the form of counts. If a data point y follows the Poisson distribution with rate *θ*, then the probability distribution of a single observation *y* is *y ∼* Poisson(*θ*). The Poisson model for data points **y_v_** = [*y*_1_*, y*_2_*, . . ., y_n_*] can be extended to the form *y_i_ ∼* Poisson(w_i_*θ*), where the *w_i_* values are known positive explanatory values proportional to the population, and *θ* is the unknown parameter of interest. Seed number in each graph can be modeled as a Poisson distribution where *y_i_* is the number of paths of some specific type (seed) in the graph, and *θ* is the underlying rate in units of seeds per graph.

To perform Bayesian inference, we need a prior distribution for the unknown rate. We use a gamma distribution as prior, which is conjugate to the Poisson. With prior distribution Gamma(*α, β*), the resulting posterior distribution is obtained as

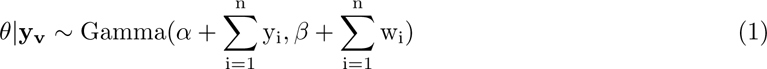

The known form of the prior and posterior densities can be used to find the marginal distribution for a single observation, which has a predictive distribution as

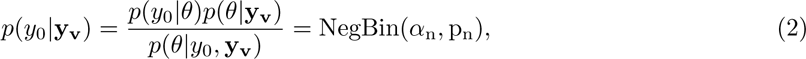

where 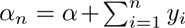, and 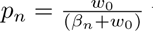 with 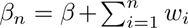. Assuming that *y*_0_ is the seed number in the primary graph, pvalue for some specific observation is obtained as

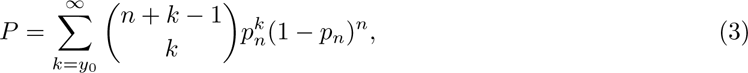

where *y*_0_, and *y_i_*, *i* = 1, 2*, . . ., n* are the number of ZNIC sites of the specific seed in primary and control data, respectively. *w_i_*numbers are the total number of ZNIC seeds in the respective graph. In our experiments, prior *α* and *β* values are assumed to be equal to *y*_0_, and *w*_0_, respectively. The first nEval significant seeds according to this criterion are passed to the next stage of refinement and enrichment.

### Refinement and Nested Seed Enrichment

The refinement and seed enrichment both use the same process of enrichment, except that refinement is only one iteration, and seed enrichment is NREFIter iterations. nEval motifs from initial evaluation step are first enriched for one iteration and top NREF motifs are further refined in seed enrichment block for NREFIter iterations or until pvalue does not improve. At each iteration, all seeds with positive likelihood ratio scores with respect to the PWM matrix obtained from the previous iteration are sorted with their score and pvalues obtained in the initial evaluation block and their ZNIC counts are computed. More specifically, it’s computed how many ZNIC samples each seed contributes to the previous samples. These counts are used to obtain significance with the same negative binomial test obtained in 4. For the negative control data, sum of the counts over nTrain control data are used for significance computation.

The PWM score that minimizes pvalue (maximizes absolute log pvalue) is selected to create the PWM matrix for the next iteration. There is an option in the algorithm to use differential enrichment where seeds are added to the PWM until the pvalue is decreasing. For example, if there are four PWM score thresholds with ZNIC log pvalues, [*−*10*, −*11*, −*9*, −*13], the default mode will consider lowest score threshold which is equivalent to log pvalue = *−*13, but differential pvalue will consider the threshold corresponding to log *−* pvalue = *−*11. The differential pvalue option will generally lead to a simpler motif structure. Default mode is used for bipolar and 3D hypothalamus dataset, and differential pvalue is used for the preoptic area of mouse hypothalamus.

Maximum likelihood estimation is used to estimate a new version of the motif PWM matrix for the next iteration. This step is iterated until the pvalue is decreasing or the maximum number of NREFIter is performed. In order to perform maximum likelihood estimation, let’s assume that we have L distinct cell types in our dataset, encoded as integer numbers from 1 to *L*. Assuming that the initial seed consists of *W* letters, *S* = *s*_1_*, s*_2_*, . . ., s_W_*, the PWM matrix, *M*, is a *L × W* matrix where *W* is the length of the searched motif. Elements of the PWM matrix in the first iteration are obtained as follows,

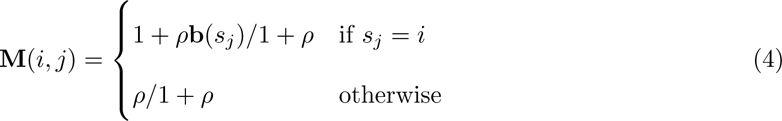

where *ρ* is the Dirichlet prior set to 0.01 in our experiment, and b is the *L ×* 1 vector of background probabilities of the cell types. For the subsequent iterations, the maximum likelihood estimation for M matrix is obtained as follows [13],

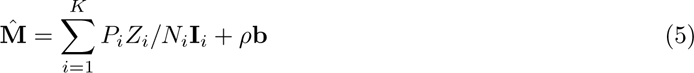

The PWM matrix, **M**, is obtained by normalizing **M̂** through the columns. *K* is the number of seeds involved in the motif, **I***_i_* is the indicator matrix of ith seed, which is equivalent to PWM matrix of 4 with *ρ* set to 0. *Zi*, *Ni*, and *Pi* are the incremental ZNIC counts, total ZNIC counts and log pvalue of the ith seed involved in the motif, respectively. Total ZNIC counts are the number ZNIC sites of the ith seed and incremental ZNIC counts are the fraction of these sites that don’t have any node in common with previous seeds, up to *i^′^*th seed.

### Applying SMORE on Synthetic and Real Data

We have tested SMORE on synthetic data and real data. The results on synthetic data helps to quantify the algorithm’s performance by assessing its ability to identify known truth patterns at various embedding frequencies. Subsequently, SMORE has been applied to multiple real spatial transcriptomic datasets, each of which is thoroughly detailed in its corresponding section.

The background frequency in synthetic data experiment is assumed to be near uniform, with

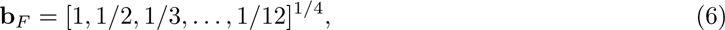

where **b***_F_* is the background frequency and it’s assumed to be known to the algorithm. Cell type labels, other than the ones for the motifs, are distributed randomly among available nodes according to their frequency. This specific form of background frequency is arbitrary and is designed to have at most twofold difference in the cell type frequency to avoid potential unwanted repeating cell type patterns like AAAA. Output motifs are represented by sequence logos. Some output motifs are simple in structure and some are more complex, consisting of multiple cell types in each position. Simple logos like AAAA indicate a repetitive pattern of A type cells interconnected in the graph within that particular tissue. In more complex motifs like (AB)AAA, the first position can be occupied by either A or B.

In 3a, the Pearson correlation coefficient (PCC) between the extracted motifs and the embedded motifs is shown. The Tomtom method [62] is employed to identify the best enriched matches among the 10 output motifs from SMORE. Tomtom searches for the best match by considering all possible shifts of the query motif with respect to the target motif (the embedded motif in this case). The matching position weight matrices (PWMs) are aligned using the obtained offsets and overlaps from Tomtom, and the PCC is computed and plotted to evaluate the pipeline’s accuracy. Each embedded motif word corresponds to a 12 *×* 4 PWM matrix, employing a Dirichlet prior with a weight of 0.01. The Dirichlet prior is a uniform distribution of the cell types.

### Computing TPR and FPR

TPR is computed as TPR = TP*/*(TP + FN), where TP (True Positive) denotes the number of output motifs with a correlation exceeding the threshold (0.95 in our context) and a pvalue below the specified threshold. FN (False Negative) is the number of instances with a correlation surpassing the threshold but a pvalue greater than the specified threshold. Thus, TP + FN represents the overall number of cases with a correlation exceeding the threshold.

FPR is defined as FPR = FP*/*(FP + TN), with FP (False Positive) indicating the number of output motifs possessing a correlation below the threshold and a pvalue below the specified threshold. TN (True Negative) is the number of output motifs with a correlation below the threshold and a pvalue exceeding the specified threshold. Consequently, FP + TN denotes the total number of cases with a correlation below the threshold.

### Gene Expression Analysis

One potential method for deriving a functional interpretation of these motifs involves examining gene expression disparities between the cells participating in the motif and those that are not involved. This analysis is carried out in section 3.4. Each motif is composed of multiple positions, and each position includes one or more cell types. For example, the motif (AB)CDE comprises cell types A and B in its first position. Among all cells labeled as A (*N_A_*), only a subset, *N_AM_*, takes part in the initial position of this particular motif. Each cell has a gene expression profile. For each gene, the delta median(dMedian) is calculated by subtracting the median expression of the gene in the subset *N_AM_* cells involved in the motif from the median expression of that gene across all *N_A_* cells of the specific type. The significance for this delta median value is then computed against a random selection of *N_AM_* cells from the total *N_A_* cells of that type.

Assume that there are *N_A_* cell types of A, *N_AM_* of these cell types is in motif (AB)CDE, with dMedian expression *x*_0_. One sided pvalue (significance) for these cells is *p*(dMedian *≥* x_0_). The probability that dMedian expression of these *N_AM_* cells is larger than *x*_0_ is the probability that at least *N*_0_ = floor(N_AM_*/*2 + 1) of these cells have expression greater than *x*_0_,

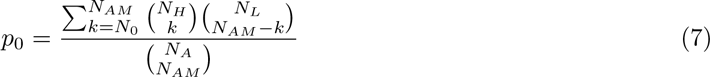

Where *N_L_* is the number of cells (out of all *N_A_* cells) that have lower than *x*_0_ delta expression, and *N_H_* is the number of cells that have higher than *x*_0_ delta expression.

---

**Figure S1:**
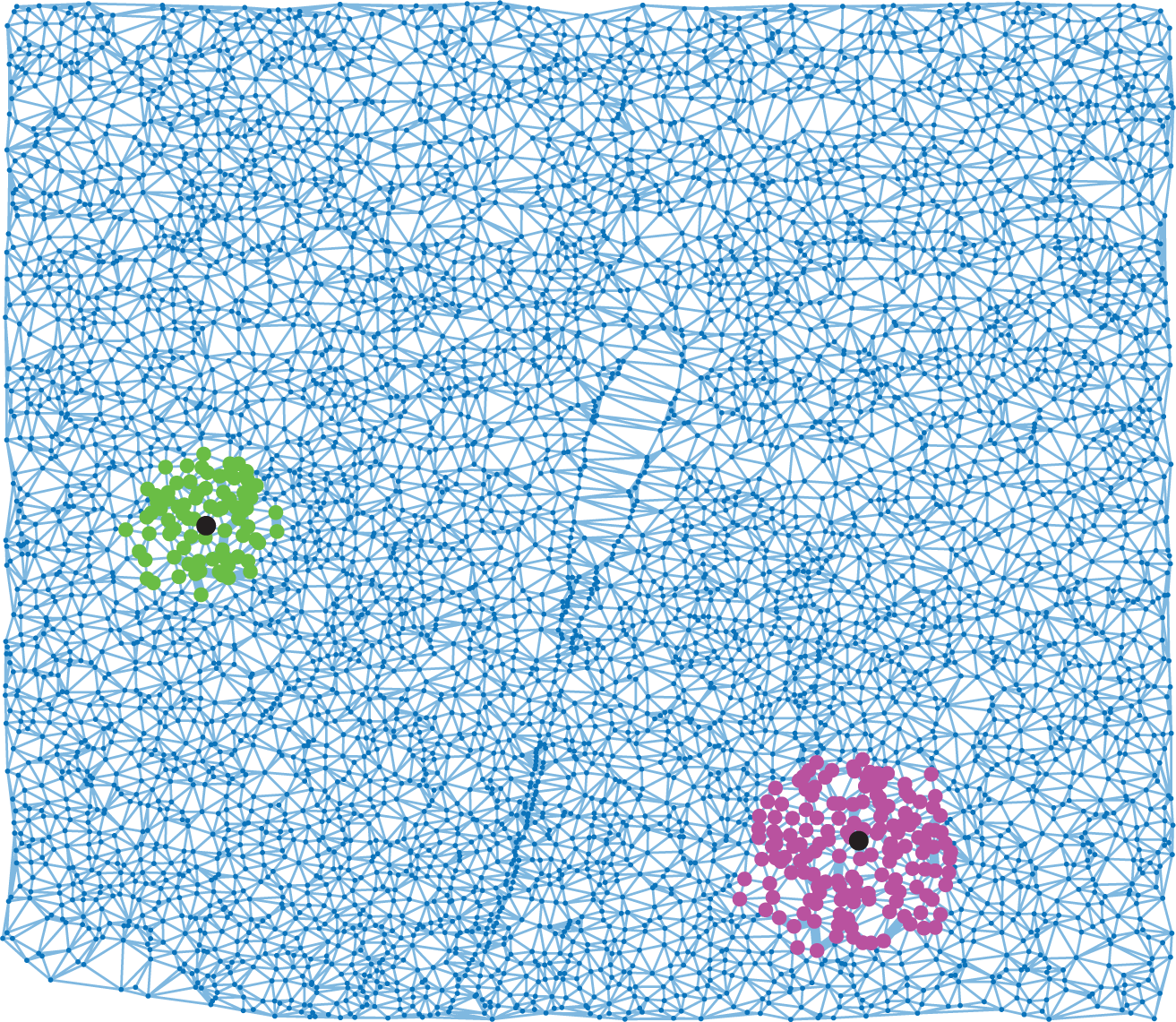
Examples of kernel neighborhoods. An example of the kernel around the center black colored node from the mouse hypothalamic preoptic region dataset. Highlighted nodes in green are length 4 kernel nodes and magenta-colored nodes are length 6 kernel nodes.

**Figure S2:**
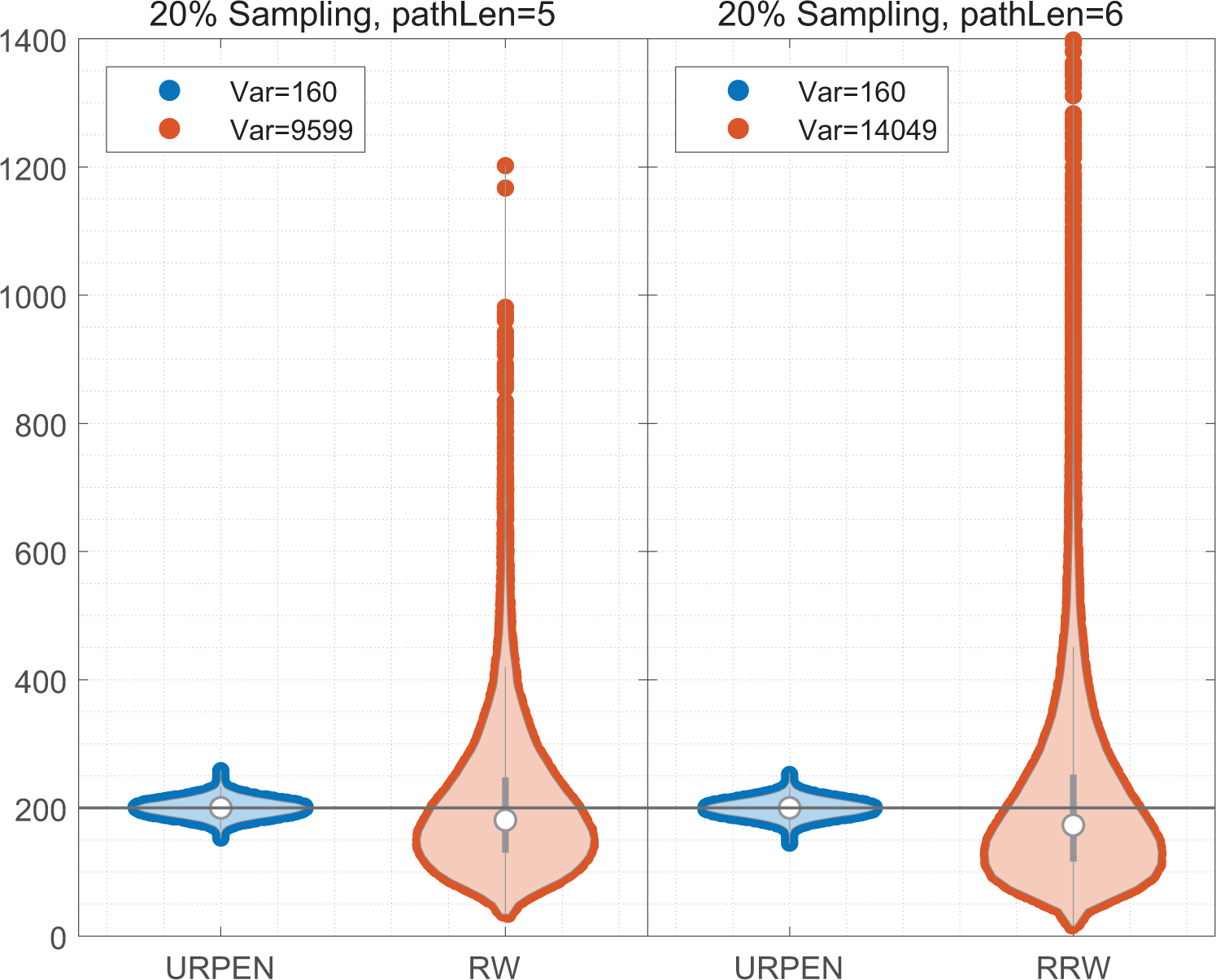
Unbiased sampling of paths from neighborhood graphs. URPEN and random walk (RW) is used to sample 20% of the graph described in Fig. 2 for length 5 non-radial samples and length 6 radial samples. In both cases, URPEN samples paths uniformly, while random walk is not uniform. The sampling probability in URPEN is set to p = (1, 1, . . ., 1, 0.2) and the test was performed 1000 times.

**Figure S3:**
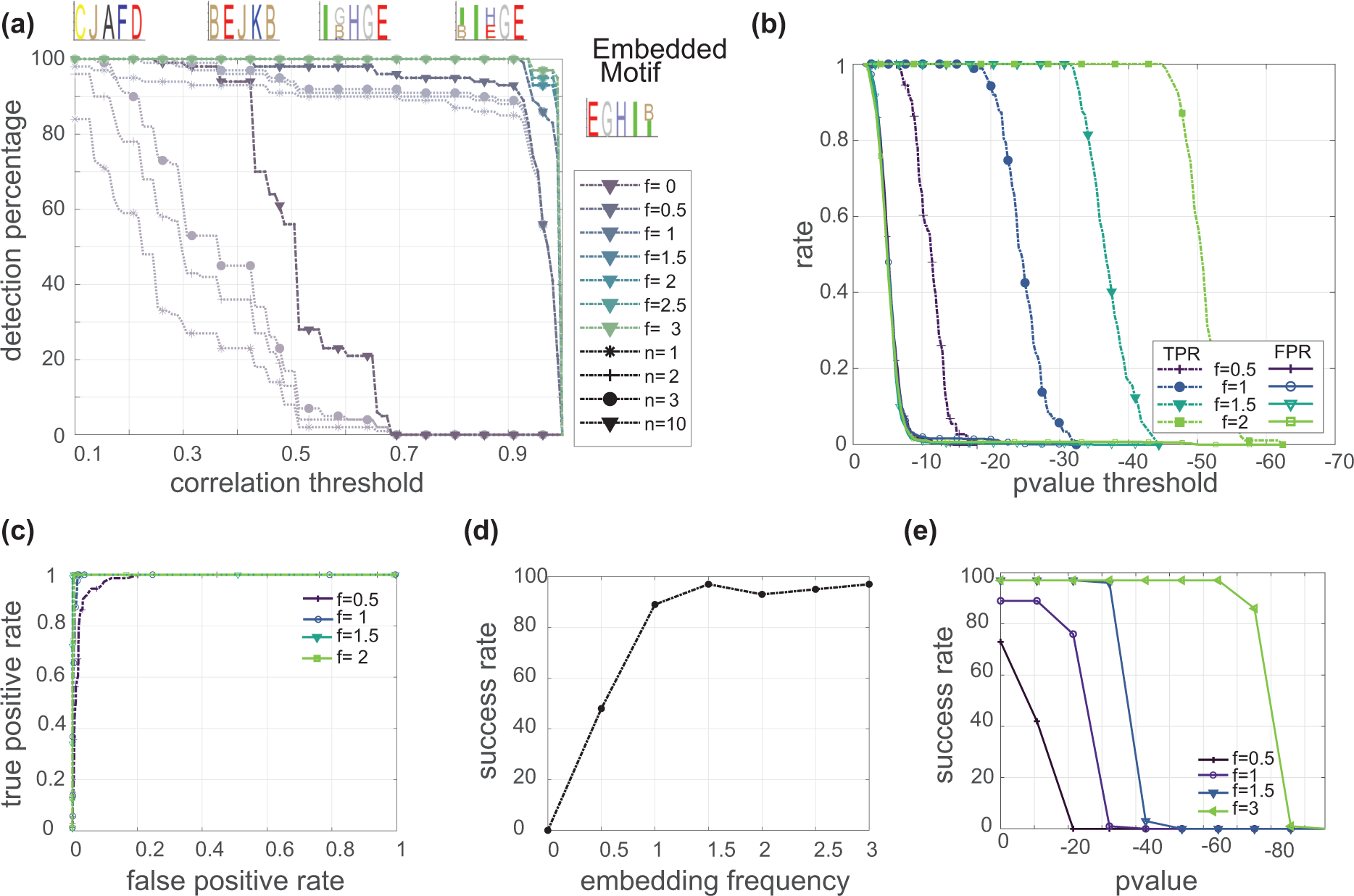
Evaluation of SMORE’s performance on synthetic data with known ground truth. Evaluation of SMORE’s performance on synthetic data with known ground truth. Same as Fig. 3, for length 5 motif.

**Figure S4:**
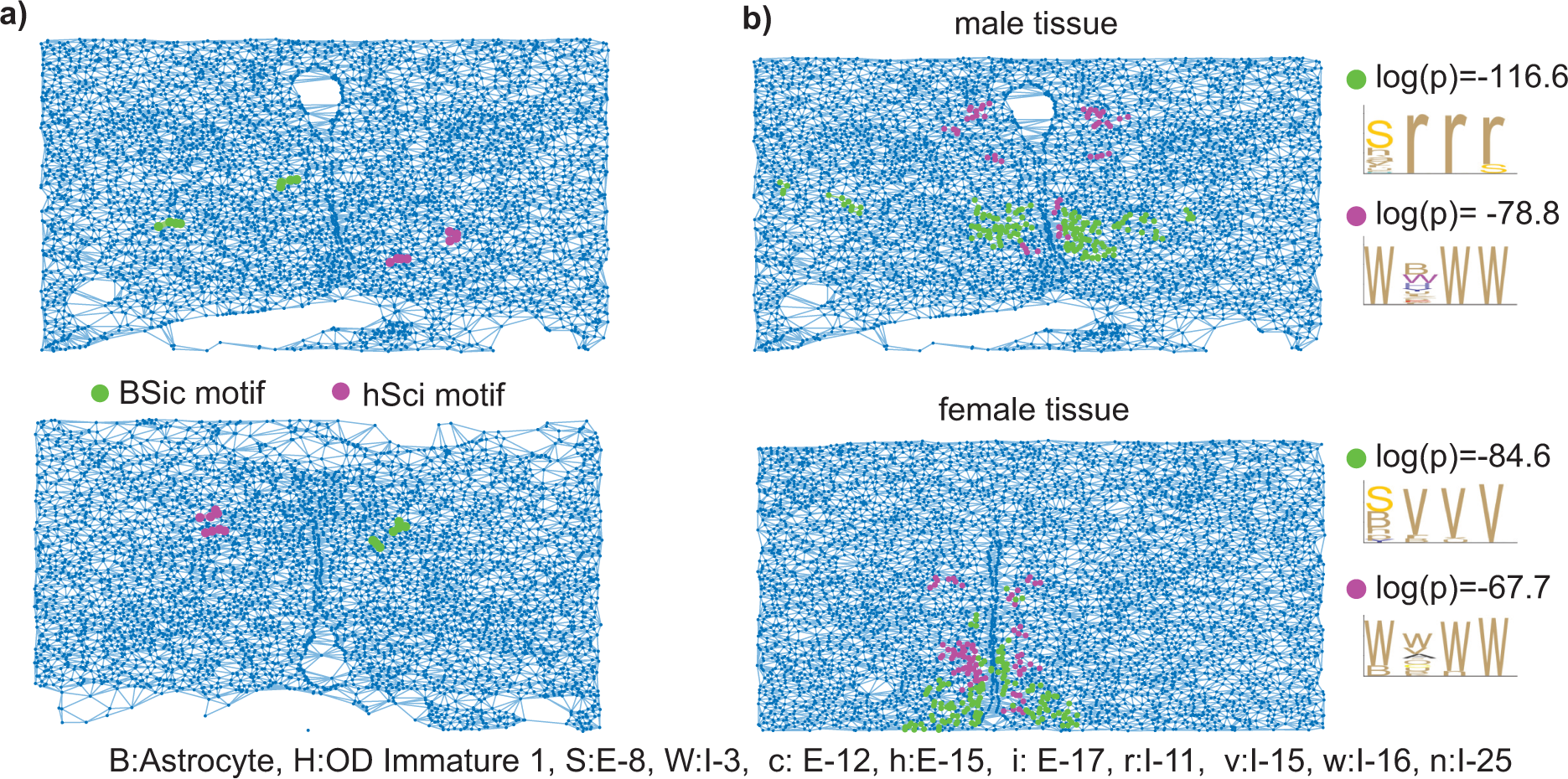
Highlights of the specific motifs in mouse hypothalamic preoptic region. **(a)** Highlight of two similar motifs in male tissues, animal IDs, 10, and 11. Involved cell types are B: Astrocyte, S: E-8, c: E-12, h: E-15, i: E-17. **(b)** Two similar motifs in male and female tissues. First motif in male tissue mostly consisted of (S: E-8, h: E-15, á: I-34) in the first position and r: I-11 in the remaining ones. For the female tissue, the first motif mostly consists of (S: E-8, h: E-15, p: E-23, Y: I-7) in the first position and v: I-15 in the remaining positions. The second motif which mostly consists of w: I-16 for both tissues, involves (W: I-3, Y: I-7, t: I-13) in the second position of the male motif and (w: I-16, v: I-15, Õ: I-10) in the second position of the female motif.

**Figure S5:**
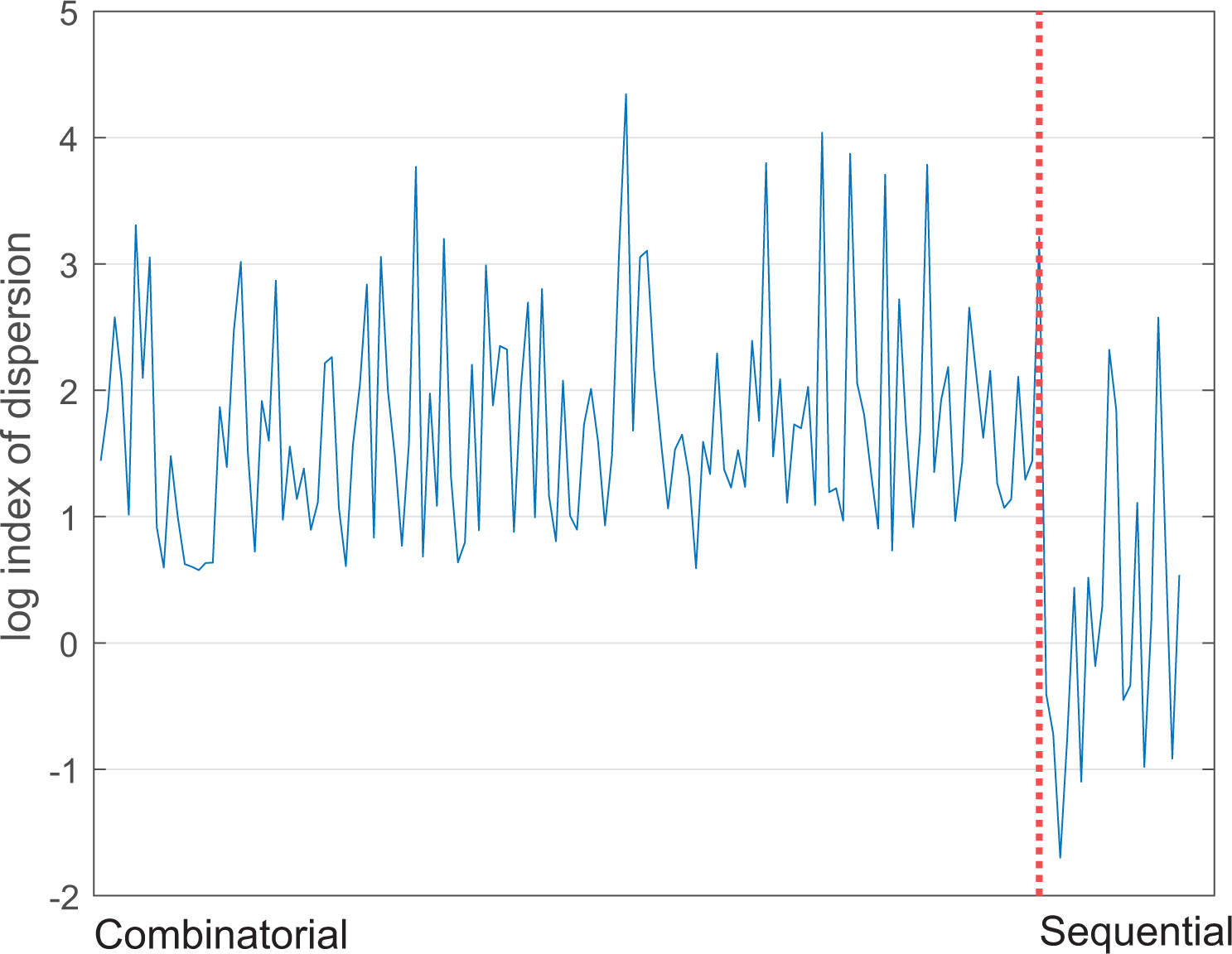
Index of dispersion for combinatorial and sequential genes. Index of dispersion for gene expression values obtained through combinatorial MERFISH and sequential smFISH. This coefficient is defined as the ratio of variance over mean. Negative log values (index of dispersion less than 1) are under dispersed. All combinatorial genes are over dispersed, while most sequential ones are under dispersed.

